# RT-Sort: an action potential propagation-based algorithm for real time spike detection and sorting with millisecond latencies

**DOI:** 10.1101/2024.04.08.588620

**Authors:** Tjitse van der Molen, Max Lim, Julian Bartram, Zhuowei Cheng, Ash Robbins, David F. Parks, Linda R. Petzold, Andreas Hierlemann, David Haussler, Paul K. Hansma, Kenneth R. Tovar, Kenneth S. Kosik

## Abstract

With the use of high density multi electrode recording devices, electrophysiological signals resulting from action potentials of individual neurons can now be reliably detected on multiple adjacent recording electrodes both *in vivo* and *in vitro*. Spike sorting assigns these signals to putative neural sources. However, until now, spike sorting can only be performed after completion of the recording, preventing true real time usage of spike sorting algorithms. Utilizing the unique propagation patterns of action potentials along axons detected as high fidelity sequential activations on adjacent electrodes, together with a convolutional neural network based spike detection algorithm, we introduce RT-Sort (Real Time Sorting), a spike sorting algorithm that enables the sorted detection of action potentials within 7.5ms±1.5ms (mean±STD) after the waveform trough while the recording remains ongoing. RT-Sort’s true real-time spike sorting capabilities enable closed loop experiments with latencies comparable to synaptic delay times. We show RT-Sort’s performance on both Multi-Electrode Arrays as well as Neuropixels probes to exemplify RT-Sort’s functionality on different types of recording hardware and electrode configurations.

## Introduction

Over the past decade, significant improvements in extracellular neural recording techniques have resulted in the development of dense multi electrode recording devices for both *in vivo* (Jun et al. 2017, Ye et al. 2023) and *in vitro* (Ballini et al. 2014) recordings. These advances have significantly enhanced our understanding of neural circuits and their activity. With interelectrode pitches smaller than 20µm, these devices allow for the recording of the activity of a single neuron by multiple adjacent electrodes. Still, in contrast to intracellular single cell recordings, individual extracellular electrodes regularly detect the activity of multiple adjacent neurons. To address this problem, spike sorting algorithms have been developed for the assignment of detected spikes to specific units that represent putative neurons (Hilgen et al. 2017, Pachitariu et al. 2018, Jun et al. 2017b, Yger et al. 2018, Garcia & Pouzat 2015, Diggelmann et al. 2018).

However, current conventional spike sorting algorithms are far from ideal (Buccino et al. 2022, Steinmetz et al. 2018). The sorting algorithms are usually based on spike waveform template matching. This means that the detected spikes are clustered into groups with similar waveform shapes, which then determines the units to which these spikes are assigned. This method does not account for time variant biological events or experimentally induced phenomena that alter the shape of the waveform. For example, extensive preceding activity of a bursty neuron can significantly change the shape of the waveform (Erickson et al. 1996, Bean 2007, Stratton 2012). This is especially problematic when studying neural activity that occurs in occasional barrages like hippocampal sharp wave ripples or population bursts of *in vitro* cultures. Furthermore, pharmacological treatments affecting the ion channel dynamics and/or differences in temperature can lead to large differences in the detected waveform shape as well (Erickson et al. 1996, Bean 2007). As a result, spikes from a single neuron can often be assigned to multiple differently detected units or even completely missed. This causes the most commonly used spike sorting algorithms to be in stark disagreement with each other (Buccino et al. 2020). Notably, this cited study from Buccino et al. showed that when six different, regularly used, spike sorting algorithms (Hilgen et al. 2017, Pachitariu et al. 2018, Jun et al. 2017b, Yger et al. 2018, Garcia & Pouzat 2015, Diggelmann et al. 2018) were applied to the same Neuropixels recording (Siegle et al. 2021), out of the total 1356 uniquely detected units, all sorting algorithms were in agreement with the detected spike times for only 33 units while 1093 units were detected by only one of any of the sorting algorithms.

Furthermore, when two neurons closely located to the same electrode fire an action potential at the same time, it will lead to an overlap of the waveforms in the recorded signal. This overlapping signal will not match any of the templates and will therefore often be missed by the sorter (Garcia et al. 2022). Finally, template matching based algorithms require all spikes in the recording to be pooled together before performing the clustering. As a result, spike sorting can only be performed after a recording is finished, leaving it impossible to know to which neuron a detected spike belongs while an experiment is ongoing.

This issue will impact anyone interested in performing closed loop experiments in which a neural system receives external inputs based on preceding activity recorded from the same neurons. Such closed loop experiments have been performed in multiple forms, ranging from controlling the bursting activity of *in vitro* cultures (Wagenaar et al. 2005) to embedding a neural culture in a virtual environment (DeMarse et al. 2001, Kagan et al. 2022) or letting a neural culture control a robot in the real world while providing information about the robot’s environment (Bakkum et al. 2007). Similar issues are faced when reading out signals of brain machine interfaces in real time (Patil et al. 2008). A more in depth review of various types of closed loop experiments and the difficulties with real time spike sorting can be found in (Arsiero et al. 2007) and (Franke et al. 2012) respectively.

As a result, various groups have attempted to develop algorithms that can sort detected action potentials within tens of milliseconds after their occurrence, in the range of signal transmission delay times of single synapses (Koch et al. 1996). However, so far no algorithm has yielded good results. For example, Franke et al. suggested a method akin to template matching, using linear filters and Independent Component Analysis (ICA) for the real time detection and sorting of action potentials (Franke et al. 2010). However, in subsequent work co-authored by Franke, it was noted that: *“The analysis of extracellular neuronal signals, recorded at high spatiotemporal resolution, reveals that the recorded data cannot be modeled as a purely linear mixture. As a consequence, ICA fails to separate completely the neuronal signals and cannot be used as a stand-alone method for spike sorting in HD-MEA recordings.”*, concluding that post processing techniques have to be used to overcome the most severe limitations of ICA (Jackel et al. 2012). This makes the ICA based method useful as a preprocessing step to spike sorting but defeats its purpose as a real time spike sorter.

More recently, machine learning based solutions have been proposed. For example, Li et al. utilized a convolution neural network (CNN) trained with manually labeled spikes to sort spikes detected on a single electrode (Li et al. 2020). They claimed that their method could classify spikes in one millisecond; however, the work did not include results of the actual latency between the occurrence of the spike and their classification which creates uncertainty about the actual sorting latencies in real time. More importantly, this method entailed training a model for each individual electrode, requiring manually labeling spikes for each electrode before being able to run a real time experiment. In addition, Li et al. reported on lower performances when only several hundred manually labeled spikes were included for the training and also noted that a spike waveform may be recorded by multiple electrodes at the same time, which still requires a solution for distinguishing spikes that are simultaneously recorded by different electrodes. Therefore, it can be concluded that although highlighting the potential of CNN’s for real time spike sorting, the algorithm developed by Li et al. cannot readily be applied to experiments using conventional recording devices containing hundreds of closely located electrodes (Jun et al. 2017, Ballini et al. 2014).

Also utilizing CNNs, Racz et al. took advantage of the fact that single spikes can be detected on multiple electrodes (Racz et al. 2020). They trained their model using data from a silicon probe, resulting in a data matrix representing the signal of 32×4 equally distanced electrodes, to better distinguish waveforms from closely located but distinct neurons. They applied Kilosort, an offline, template matching based spike sorting algorithm (Pachitariu et al. 2016) to a pre-recording made by the probe to obtain a “ground truth” for training their model. As a result, a new model would have to be trained every time a new probe was inserted. Furthermore, by optimizing the models to an electrode configuration of 32×4 electrodes, this algorithm doesn’t transfer well to other related recording devices with variable electrode configurations (Jun et al. 2017, Ballini et al. 2014). Most importantly, the authors noted that the model was directly trained on the outputs of Kilosort and so any mistakes resulting from the previously listed shortcomings of template matching based sorting algorithms are necessarily inherited by this algorithm.

We set out to develop RT-Sort (Real Time Sorting), a spike detection and sorting algorithm that overcomes the issues related to overlapping spikes, changes in waveform shapes and the inability to perform spike sorting in real time. Instead of relying on waveform shape similarity, we based our sorting method on a more biologically plausible phenomenon. Namely, the fact that an action potential propagates along a neural axon and thus gets detected with very high fidelity by a sequence of electrodes, in the same order, with sub millisecond but nonzero time delays (Tovar et al. 2018). Interestingly, all real time spike sorting methods described above still apply some form of template matching, where the shape of the spike waveform recorded on one or multiple electrodes directly determines the unit cluster to which the spike is assigned. For the first time, we used action potential propagation detection instead of spike waveform template matching to identify individual neurons in our recordings. This idea has already been acknowledged and utilized in the classification of neuron types based on high density recordings (Jia et al. 2019) but has not yet explicitly been applied to enhance spike sorting routines.

In this work, we present RT-Sort, an algorithm that can sort spikes in real time based on their action potential propagation trajectory. In doing so, we show that RT-Sort performs at the level of a broad series of conventionally used template matching based spike sorting algorithms (Hilgen et al. 2017, Pachitariu et al. 2018, Jun et al. 2017b, Yger et al. 2018, Garcia & Pouzat 2015, Diggelmann et al. 2018), while reporting latencies for detection and sorting of 7.5ms±1.5ms (mean±STD) from the trough of the waveform on replayed recordings. Furthermore, RT-Sort can be applied to any system with sufficiently high electrode density (≤50µm pitch) as exemplified by our implementations on some of the most commonly used recording devices for both *in vivo* (Jun et al. 2017) and *in vitro* (Ballini et al. 2014) experiments.

Because our method relies on the sequential detection of a rapidly decaying propagating action potential on multiple electrodes, our spike detection sensitivity is an important limiting factor as to whether we can sort a spike. Relying on conventional RMS based threshold crossings does not provide the desired signal to noise ratio. Therefore, similar to Racz et al., we utilize the power of CNNs for the detection of spikes. However, thanks to our methods of artificial data amplification, inspired by the work from Parks et al. (Parks et al. 2018), our model doesn’t require training on individual recording or electrode configurations. Furthermore, it allows us to generate very large training samples, which enables the development of a more powerful and generalizable model.

As a result, a single model trained for a specific model type – recording device combination can be applied to any future experiment on that model with the specific recording device. We have trained readily available models for *in vitro* human brain organoid neurons recorded with Maxwell Biosystems HD-MEAs and *in vivo* mouse neurons recorded with Neuropixels. We will also release a straightforward library to the community that allows for training other models and to applying RT-Sort under various experimental conditions.

## Results

### Real time spike detection on injected spikes

A CNN with 4 layers (Fig. 1A) was trained to detect spikes in recording device specific noise. Separate models were trained for Maxwell MEA recordings performed on human brain organoids and Neuropixels 1.0 recordings performed on mice (Sup. Fig. 1 for Neuropixels). The training was performed using averaged waveform shapes of electrodes selected along the whole recorded axon, the trajectory along which the action potential propagates (Fig. 1B). These averaged waveform shapes were then pasted into recording device specific noise fragments which enabled artificial amplification of the training dataset (Fig. 1C) and the generation of training cases of overlapping waveform shapes with ground truth (Fig. 1D). The small differences between the training loss and testing loss indicated that our models are not overfitting nor underfitting the data (Fig. 1E). See the corresponding sections in *Methods* for a more detailed description of the model training and evaluation.

**Figure 1:**
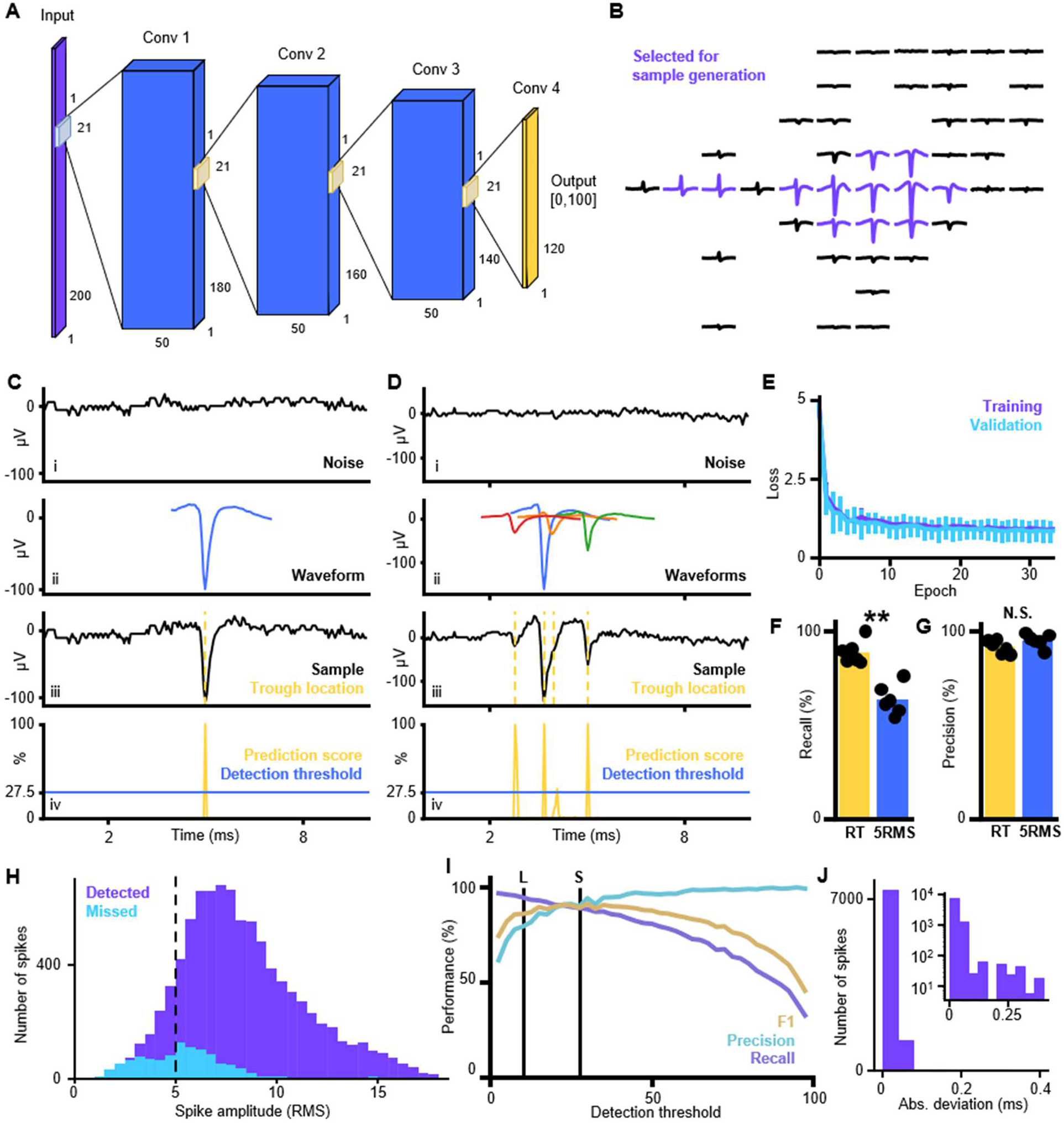
Action potential detection with high temporal accuracy using a convolutional neural network (CNN). A) Architecture of the CNN used for spike detection in 20kHz MEA recordings. The input layer consists of 200 nodes, corresponding to a 10ms window. The output layer consists of 120 nodes corresponding to 2-8ms in the input window. Each output node provides a score between 0 and 100 indicating the likelihood that the corresponding input frame contains a waveform trough. The 4 convolutional layers have a kernel size of 21 frames and a stride of 1 frame so that each output node makes a prediction based on the signal in 2ms before until 2ms after the corresponding frame in the input. B) Example of averaged waveform footprint detected by Kilosort2. The purple traces were selected for training/validating the CNN. Note the variety in the selected waveform shapes from this single unit. This ensures that the detection model can detect the propagating action potential along different parts of the neuron. C) Example of training/validating sample creation and model prediction. i: A 10ms sample of recording specific noise is taken. ii: A waveform shape is selected from the training or validating pool. iii: The waveform is pasted into the recording device specific noise with ground truth certainty about the trough location (marked with yellow dotted line). iv: CNN detection model predictions for the frames in 2-8ms of the input window show a narrow detection peak at the waveform trough. All figures share the same x-axis. D) Same as C but with multiple overlapping waveforms in the same sample. E) Training (purple) and validating (cyan) loss as a function of training epoch. Error bars indicate the STD over the different cross-validation folds. The small differences between the training loss and validating loss indicate that the models is not overfitting nor underfitting the data. F) Recall when validating the detection model on samples generated from the held-out recording and when applying a 5RMS threshold to the same samples. The markers indicate the results for each of the 6 held-out datasets and the bar reflects the mean over all held-out datasets. Mean±STD for CNN = 89.1%±5.59% and for 5RMS = 63.6%±7.36%. The detection model has a significantly higher recall (P = 1.25*10^-7^, two-sided paired t-test, n=6). G) Precision when validating the detection model on samples generated from the held-out recording and when applying a 5RMS threshold to the same samples. The markers indicate the results for each of the 6 held-out datasets and the bar reflects the mean over all held-out datasets. Mean±STD for CNN = 91.5%±3.23% and for 5RMS = 95.0%±3.36%. The difference in precision is not significant (P = 0.13, two-sided paired t-test, n=6). H) Amplitude distribution of detected (purple) and missed (cyan) spikes by the detection model shows a good detection performance for spikes below 5RMS. Amplitudes are expressed as RMS relative to the surrounding 50ms of signal. I) F1 score, precision and recall of the detection model as a function of the detection threshold. The loose and stringent detection thresholds are marked with L and S respectively. J) The absolute deviation between the model detections and the ground truth trough times shows a high temporal accuracy with on average a deviation of 13μs (0.26 frames). Inset contains the same distribution on a log scale.

When testing the trained models on the artificially generated data from the held-out organoid recordings, using the detection threshold yielding an optimal F1 score, we obtained a recall score of 89.1%±5.59% and a precision score of 91.5%±3.23% (mean±STD, Fig. 1FG). Similarly, the mouse Neuropixels recordings yielded a recall score of 84.7%±5.81% and a precision score of 86.4%±7.90% (mean±STD, Sup. Fig. 1FG). In both cases, the obtained recall was significantly larger than applying a 5RMS threshold to the injected spikes (organoids: 5RMS recall mean±STD = 63.6%±7.36%, two-sided paired t-test P=1.25*10^-7^, n=6, Fig. 1F; mouse: 5RMS recall mean±STD = 65.1%±3.37%, two-sided paired t-test P=5.19*10^-5^, n=6, Sup. Fig. 1F). Meanwhile, the precision was not significantly different between the CNN detection model and the 5RMS threshold (organoids: 5RMS precision mean±STD = 95.0%±3.36%, two-sided paired t-test P=0.13, n=6, Fig. 1F; mouse: mean±STD = 79.7%±4.60% (smaller than RT-Sort CNN model), two-sided paired t-test P=0.15, n=6, Sup. Fig. 1G). The significantly higher recall scores were due to the fact that the model could detect spikes with amplitudes below 5RMS, with the 5th percentile of the amplitude distribution over all detected waveforms at 4.08RMS for the MEA model and 3.67RMS for the Neuropixels model. Meanwhile, the models hardly missed spikes with amplitudes above 5RMS (Fig. 1H, Sup. Fig. 1H).

Adjusting the detection threshold for the organoid model from its optimal value allowed for enhanced precision scores at the cost of a slightly lower recall score. This enabled the use of both a stringent detection score at the optimal F1 score, obtained with a detection threshold of 27.5%, as well as a loose detection threshold at 10% (Fig. 1I, 17.5% and 7.5% for the Neuropixels model, Sup. Fig. 1I). Even though we considered a detection to be a true positive if it was detected within 400μs of the actual waveform trough, the average absolute deviation from the correct trough time was only 13μs and 14μs for the organoid and mouse recordings respectively (Fig. 1J, Sup. Fig. 1J), reflecting the high temporal sensitivity of the model detections. When probing the model to update an injected piece of noise signal until it reaches an optimal detection threshold value, a classical waveform shape emerges (Sup. Fig. 2) which further highlights that the model indeed is sensitive to waveform shapes in noisy signals.

### Real time spike detection on replayed recordings

We subsequently tested our model on actual spikes detected by Kilosort2 in the held-out recordings. When only considering the largest waveform amplitude electrodes, we observed precision and recall scores of 26.0%±29.4% and 83.1%±23.4% for the MEA model applied to organoid recordings and 7.47%±12.7% and 77.6%±25.8% for the Neuropixels model applied to intact mouse brain recordings (mean±STD, Sup. Fig. 3AB). When interpreting these precision scores, it has to be considered that we are comparing the spike detection model against sorted Kilosort2 detections. A sorted detection only gets detected on a single electrode while our spike detection model can detect a spike on multiple electrodes. As such, detections from nearby units might wrongfully be considered false positive spikes leading to low precision scores. In addition, Kilosort2 is no ground truth and might miss spikes or falsely detect spikes due to various possible reasons like overlapping waveforms, noise, low amplitude or waveform shape changes. This all will negatively impact the obtained performance scores.

When we considered the 4^th^ largest waveform amplitude electrodes per unit, the precision and recall were 10.6%±20.6% and 25.3%±24.5% for the MEA model applied to organoid recordings and 5.61%±10.8% and 53.1%±33.0% for the Neuropixels model applied to intact mouse brain recordings (mean±STD, Sup. Fig. 3AB). The reduced recall reflects the fact that some of the detected units will not be detectable on 4 individual recording electrodes, even though this is a requirement for spikes to be sorted by RT-Sort. However, the bimodality of the distribution of recall scores per unit indicates that there is a subset of units that can reliably get detected (Sup. Fig. 3CD) on at least 4 individual electrodes which makes them strong candidates for detection by RT-Sort.

### Real time spike sorting on ground truth datasets

Subsequently, we utilized the CNN outputs to perform spike sorting in real time using two different ground truth datasets. A detailed explanation of the spike sorting algorithm can be found in *Methods: Propagation sequence detection* and following sections. In summary, within a short pre-recording made right before the experiment, recurring sequential detections on multiple electrodes with short but non-zero temporal interelectrode intervals were detected, reflecting the propagation of an action potential. We refer to these detections as propagation signals. Detections were based on high fidelity interelectrode intervals and compared to the highest amplitude electrode in the sequence. Sequences that were highly similar in their propagation sequence were merged into single propagation signals. This then yields the final selection of propagation signal sequences that were detected. In the subsequent experiment, sequential detections were compared to the pre-detected propagation signal sequences and if there was sufficient overlap, the sequential detection was counted as a spike for the propagation signal sequence.

The first ground truth dataset consisted of recordings that were made of rat primary neurons plated on Maxwell HD-MEAs while one of the neurons was simultaneously recorded by patch clamp *(Methods: Patch-MEA ground truth recording)*. Applying RT-Sort to this recording yielded the detection of 49 different units, including the neuron that was recorded using patch clamp (Fig. 2A-C).

**Figure 2:**
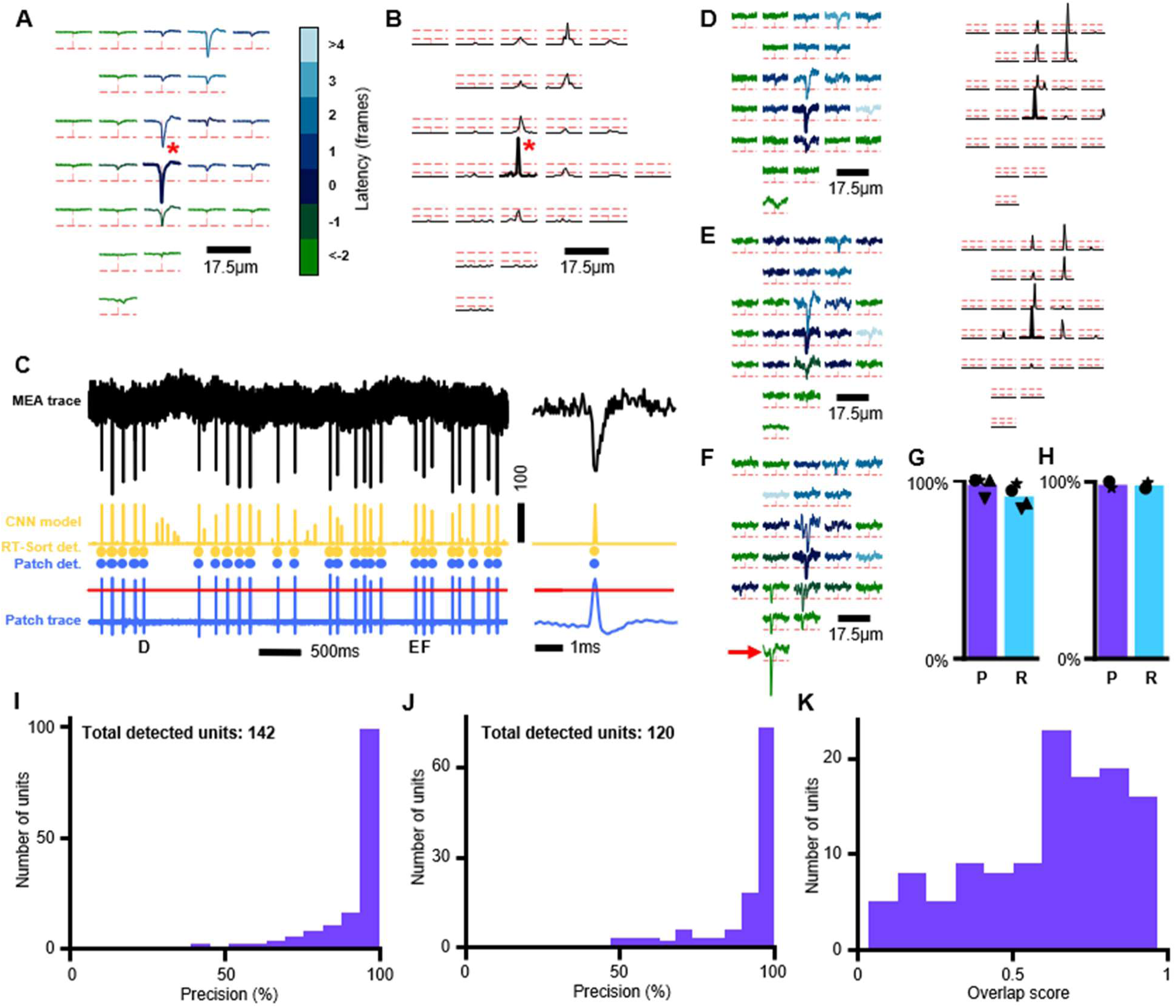
RT-Sort spike sorting performance on ground truth datasets. A) Averaged waveform footprint from a neuron recorded using a high-density multi-electrode array (MEA). Each trace represents the signal from a single electrode averaged over all action potential detections of the recorded neuron. 5 times the signal to noise ratio of the signal measured on each electrode is indicated with dotted red lines. The root electrode detected by RT-Sort and used for the spike triggered averaging is marked with the red star. The trace on each electrode ranges from 2ms before until 2ms after the waveform trough on the root electrode. All detected loose electrodes by RT-Sort are marked in bold. For each electrode, the color of the trace represents the average detection latency relative to the root electrode. B) The average spike detection model scores over all detected action potentials for the same neuron and electrodes as A. The loose and stringent detection threshold on each electrode is indicated with dotted red lines. All detected loose electrodes by RT-Sort are marked in bold. To better reflect the high temporal precision of the detection model, the time range for each trace is magnified 4 times relative to the traces in A, resulting in the detection score trace ranging from 0.5ms before until 0.5ms after the waveform trough on the root electrode (marked with red star). C) From top to bottom: Raw unfiltered MEA trace of the root electrode marked in A and B for part of the recording (black). This signal is used as input for the detection model. Detection model scores for the same period (yellow). RT-Sort spike sorted detections (yellow dots). Spike detections in simultaneously made patch recording of the same neuron (blue dots). Patch trace recorded in cell-attach mode (blue line) with spike detection threshold marked in red. Note that the detection model sometimes detects spikes from an adjacent neuron (see panel F) although RT-Sort correctly assigns these detections to a different neuron. Right: zoomed in signals for a single spike marked with D. D) Same as A and B but for a single spike from the same neuron, detected by RT-Sort in online mode. The spike is marked with D in panel C. The colors for each trace correspond to the same latencies as A. E) Another example of a single spike footprint from the same neuron, marked with E in panel C. Note the difference in the waveform shape compared to D on the electrode above the root electrode. F) Single spike footprint from the same neuron that overlaps with a spike from a different neuron (root electrode marked with red arrow). Despite the waveform overlap, the spike is still correctly detected as marked in panel C. G) Precision and recall for spikes detected by RT-Sort from 4 different neurons on 2 different MEAs with simultaneous patch-clamp ground truth recording. Precision = 97.5%±4.4%, Recall = 90.9%±5.57% (mean±STD). The example neuron in A is marked with the circle. H) Precision and recall for spikes detected by RT-Sort from 2 different neurons on an MEA with simultaneous patch-clamp ground truth recording. The detections were made using sequences detected in a pre-recording, similar to how RT-Sort would be applied in real time. Precision = 98.2%±1.78%, Recall = 97.8%±1.60% (mean±STD). I) For each unit detected in the simulated ground truth recording using sequence metrics generated based on the ground truth spike locations, the precision over all detected spikes compared to the most similar ground truth neuron. Mean±STD = 98.8%±12.4%. J) For each unit detected in the simulated ground truth recording, the precision over all detected spikes compared to the most similar ground truth neuron. Mean±STD = 97.3%±13.4%. K) For each unit detected in the simulated ground truth recording, the overlap score for the most similar ground truth neuron. Mean±STD = 0.651±0.242.

Because our CNN model was trained to detect general waveform shapes in sample specific noise which are then sorted based on their action potential propagation patterns, RT-Sort is well capable of detecting all spikes from the same neuron despite changes in waveform shape (Fig. 2DE, Sup. Fig. 4 for additional MEA example and Neuropixels example). Furthermore, since our CNN was trained on multiple waveforms injected into the same detection window, we find that RT-Sort is very suited to detect overlapping waveforms in both the injected (Fig. 1D) and actual recordings (Fig. 2F, Sup. Fig. 5 for Neuropixels examples and additional MEA examples). Finally, with the enhanced sensitivity of our spike detections, RT-Sort can detect units that are fully below 5RMS (Sup. Fig. 6). Doing the ground truth tests on n=4 neurons from 2 different MEAs using RT-Sort in offline mode, resulted in a precision of 97.5%±4.4% and a recall of 90.9%±5.57% (mean±STD, Fig. 2G). For the array with a pre-recording, RT-Sort was also applied in online mode on the n=2 neurons on this array, yielding a precision of 98.2%±1.78% and a recall of 97.8%±1.60% (mean±STD, Fig. 2H).

We further tested RT-Sort on a different ground truth dataset consisting of a simulated mouse *in vivo* Neuropixels dataset used in (Buccino et al. 2020). This simulated dataset consisted of 250 biophysically detailed neurons that each exhibit independent Poisson firing of spikes with added Gaussian noise. To first test the ability of RT-Sort to assign spikes to the correct sequences, sequence metrics were initially obtained using the ground truth spike times from the modeled recording. Subsequently assigning spikes to these sequences yielded 142 units that were detected on the minimally required 4 electrodes to measure the propagating action potential. Over these 142 units, a precision of 98.8%±12.4% (mean±STD) was obtained, reflecting a low false positive detection rate on spike detection (Fig. 2I). Subsequently, RT-Sort was run in offline mode on the first 5 minutes of the simulated dataset to detect propagation sequences, yielding 120 detectable action potential propagation sequences. Over these 120 units, a precision of 97.3%±13.4% (mean±STD) was obtained, again reflecting a low false positive rate (Fig. 2J), in accordance with the patch-MEA results.

These results indicate that RT-Sort is capable of detecting a subset of units with low false positive rates while being unable to detect other units. This is likely due to the relatively stringent detection constraints that are imposed on RT-Sort, in order to reliably detect propagating action potentials. Unlike the patch-MEA recordings, RT-Sort did miss some of the true positive spikes in the simulated dataset (Sup. Fig. 7AB). This resulted in an overlap score per unit of 0.65±0.24 (mean±STD, Fig. 2K). The missed true positive spikes could partially be due to the fact that the spike detection model was trained on real waveforms while the biophysically detailed neurons did not always yield realistically looking waveforms (Sup. Fig. 7C). RT-Sort is more sensitive to not detecting unrealistically looking waveform shapes than template matching algorithms, which makes it less likely for RT-Sort to detect false positive spikes.

### Detection latencies

For RT-Sort to be used in real time experiments, it is important to minimize the latency between the occurrence of the action potential and its detection. In our case, the latency between the waveform trough and its sorted detection can be ascribed to three different factors: biological constraints, detection speed and sorting speed (Fig. 3A). The biological constraints consist of the duration of the remainder of the spike waveform and propagation to occur after the waveform trough. For the spike detection, using a single NVIDIA-RTX-A5000 GPU allowed us to perform a forward pass through the Maxwell MEA spike detection model for 1020 electrodes simultaneously (total number of available recording electrodes for Maxwell Biosystems HD-MEAs) in 1.12ms and a forward pass through the Neuropixels spike detection model for 384 electrodes simultaneously (total number of available recording electrodes on a Neuropixels probe) in 0.92ms. Adding additional electrodes from there only led to a linear increase in the required computation times (Fig. 3B). A first order linear regression model fitted to the forward pass duration yielded: duration (ms) = 0.155 + 0.001 * #elec (R^2^=0.997, P=4.14*10^-15^) for the MEA model and duration (ms) = 0.155 + 0.002 * #elec (R^2^=0.999, P=4.09*10^-19^) for the Neuropixels model. The steeper slope for the Neuropixels model is due to the higher sampling rate of 30kHz compared to 20kHz.

**Figure 3:**
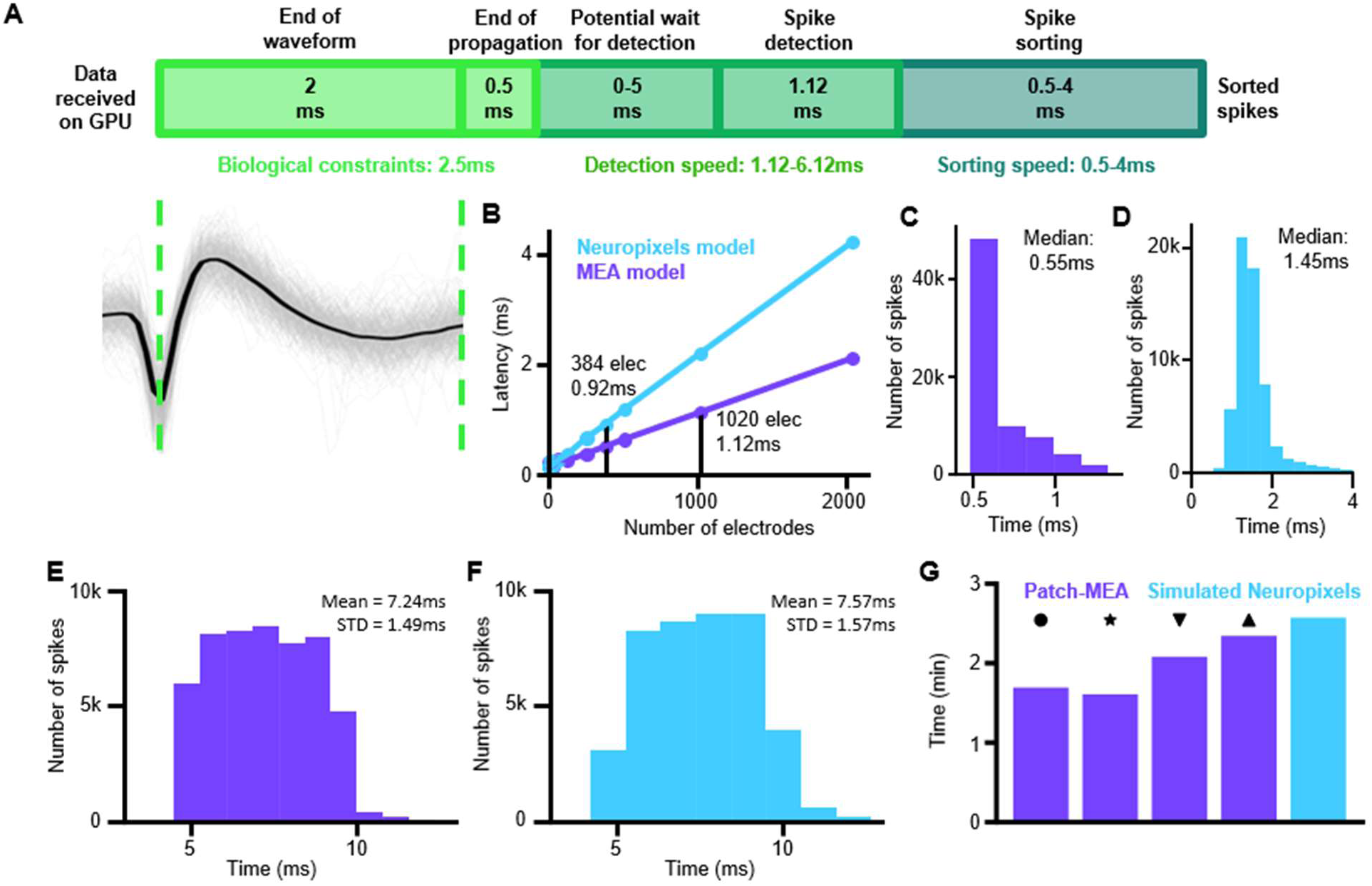
Real time spike sorting with latencies in the range of synaptic transmission. A) Schematic representation of the durations of the different steps to go from action potential trough to sorted spike. The duration is split up into 3 categories: Biological constraints refer to biologically intrinsic waiting times since the action potential trough occurs at the soma in order to measure all the data required for the detection. This consists of measuring the action potential until the end of the waveform (bottom left) plus an additional 0.5ms for the action potential to propagate. Detection speed refers to the duration for the spike detection model to detect spikes on all the electrodes used in the recording. Sorting speed refers to the duration to take the spike detection probability outputs and use those to assign spikes to the correct units. B) Computation time to perform a forward pass through the spike detection model as a function of the number of electrodes in the recording. A first order linear regression model fitted to the forward pass duration yielded: duration = 0.155 + 0.001 * #elec (R2=0.997, P=4.14*10^-15^) for the MEA model in purple and duration = 0.155 + 0.002 * #elec (R2=0.999, P=4.09*10^-19^) for the Neuropixels model in cyan. C) Distribution of spike sorting duration on the detection model outputs for the two patch-MEA with real time sorting from Figure 2H. D) Distribution of spike sorting duration on the detection model outputs for the real time detections on the simulated Neuropixels recording from Figure 2I-K. E) Distribution of spike detection and sorting durations for the real time sorting on the two patch-MEA recordings from Figure 2H. Time reflects duration from waveform trough until sorted detection (mean±STD over all detected spikes: 7.24ms±1.49ms). F) Distribution of spike detection and sorting durations for the real time sorting on the simulated ground truth dataset from figure 2I-K. Time reflects duration from waveform trough until sorted detection (mean±STD over all detected spikes: 7.57ms±1.57ms). G) Durations for detecting propagation sequences and sorting all spikes in the different offline patch-MEA recordings from Figure 2G (purple, markers same as Figure 2G) and in the pre-recording for the simulated Neuropixels recording made before running RT-Sort in online mode for Figure 2I-K (cyan). Time is expressed as number of minutes per minute of recording time.

Based on these results, we conclude that our model is very well suited for detecting action potentials within milliseconds of their occurrence, at a higher sensitivity than thresholding the signal and all while being robust against overlapping waveforms as well as changing waveform shapes.

Subsequently processing the model detections into sorted spikes only required an additional 0.5-4ms with a distribution median of 0.55ms (Fig. 3C) for the patch-MEA recordings from Fig. 2H and a distribution median of 1.55ms (Fig. 3D) for the simulated Neuropixels recording from Fig. 2I-K. Altogether, this yielded a total latency from waveform trough to sorted detection of 7.24ms±1.49ms (Fig. 3E) for the patch-MEA recordings and 7.57ms±1.57ms (Fig. 3F) for the simulated Neuropixels recording (mean±STD) when applying RT-Sort in online mode. These results clearly illustrate the detection and sorting speed enabled by RT-Sort. Running RT-Sort in online mode requires detecting sequences in offline mode first in a pre-recording. Detecting sequences in offline mode took 1.93±0.30 minutes (mean±STD) per minute of recording for the patch-MEA recordings and 2.58 minutes per minute of recording for the simulated Neuropixels recording (Fig. 3G). Therefore, RT-Sort can be used in online mode with sequences recorded only a few minutes in advance.

### RT-Sort compared to other sorters

We next compared RT-Sort to 6 different conventionally used sorting algorithms (SpyKING Circus, HDSort, HerdingSpikes2, IronClust, Kilosort2, IronClust and Tridesclous (Hilgen et al. 2017, Pachitariu et al. 2018, Jun et al. 2017b, Yger et al. 2018, Garcia & Pouzat 2015, Diggelmann et al. 2018)) that were all applied to the same mouse Neuropixels recording (Siegle et al. 2021). RT-Sort detected a total of 168 units in the Neuropixel recording, which is fewer than any of the other sorters, in agreement with the results observed in the simulated dataset (Sup. Fig. 8A). The units consisted of detected propagations of 40µm to 150µm in length (Sup. Fig. 8B) and up to 500µs in duration (Sup. Fig. 8C).

We sought to quantify the quality of the detected units by comparing them to the other sorters. A previous comparison between the six other sorters on this dataset showed only a very limited overlap in the detected units between the different sorters (Buccino et al. 2020) (see *Methods: Overlap score* for how the overlap was assessed). We hypothesize that the low overlap between units could be due to false positive spike detections of different sorters which could make the spike trains of different sorters so different that they are not recognized as coming from the same unit. This would mean that good performance of a certain sorter could potentially be obscured by bad performance of a different sorter (Fig. 4A). As a result, for comparing detections between sorters, we quantified how similar the spikes that were detected by either RT-Sort or a comparison sorter alone are compared to the spikes that both sorters agreed on.

**Figure 4:**
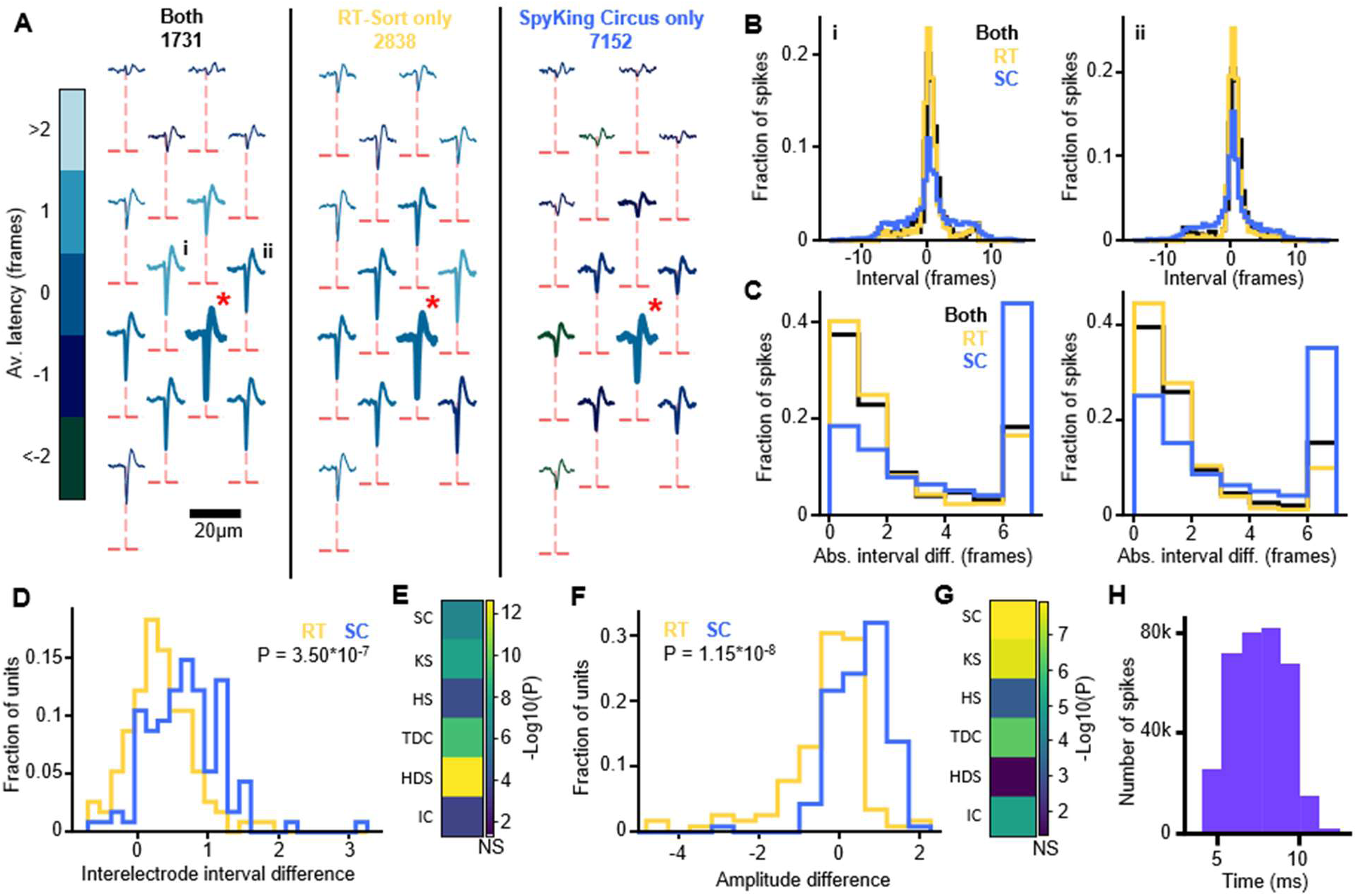
RT-Sort detects spikes with consistent propagations and amplitudes. A) Waveform footprint of a unit detected by both RT-Sort and SpyKing Cricus. Left: averaged footprint over all spikes detected by both RT-Sort and SpyKing Circus. Middle: averaged footprint over all spikes only detected by RT-Sort. Right: averaged footprint over all spikes only detected by SpyKing Cricus. 5 times the signal to noise ratio of the signal measured on each electrode is indicated with dotted red lines. The root electrode detected by RT-Sort and used for the spike triggered averaging is marked with the red star. All detected loose electrodes by RT-Sort are marked in bold. For each electrode, the color of the trace represents the average detection latency relative to the root electrode. B) The distribution of interelectrode intervals per spike relative to the root electrode for the comparison electrodes marked in A. Each spike group in A (both, RT-Sort only and SpyKing Circus only) is plotted as a separate distribution. The wider base of the SpyKing Circus distribution indicates spike contamination from different units for the SpyKing Circus only spikes, which is not detected by RT-Sort C) The distribution of interelectrode interval differences compared to the mean of all spikes detected by both RT-Sort and SpyKing Circus for the comparison electrodes in B. Differences are clipped at 7 frames. D) Interelectrode interval difference scores for all matching RT-Sort and SpyKing Circus units. SpyKing Circus only spikes are significantly more different from the matched spikes in their interelectrode intervals compared to the RT-Sort only spikes (P = 3.50*10^-7^, one-sided paired t-test). E) −Log10(P) values for inter-electrode interval differences comparing RT-Sort only spikes to other sorter only spikes relative to matched spikes. All other sorters are significantly more different (P<0.05, one-sided paired t-test). Abbreviations: SC = SpyKing Circus, KS = Kilosort2, HS = Herdingspikes2, TDC = Tridesclous, HDS = HD-Sort, IC = IronClust. F) Amplitude difference scores for all overlapping RT-Sort and SpyKing Circus units. SpyKing Circus only spikes are significantly more different from the matched spikes in their amplitudes compared to the RT-Sort only spikes (P = 1.15*10^-8^, one-sided paired t-test). G) −Log10(P) values for amplitude differences comparing RT-Sort only spikes to other sorter only spikes relative to matched spikes. All other sorters are significantly more different (P<0.05, one-sided paired t-test) except HD-Sort (P=0.0789, one-sided paired t-test). Abbreviations same as E. H) Distribution of spike detection and sorting durations for the real time sorting on the Neuropixels dataset from figure 4. Time reflects duration from waveform trough until sorted detection (mean±STD over all detected spikes: 7.62ms±1.58ms).

Firstly, the distribution of the interelectrode intervals between each loose electrode in the sequence and the root electrode was constructed for “matched” spikes, “RT-Sort only” spikes and “Other sorter only” spikes (Fig. 4B). For every spike in each of these three groups, the interelectrode interval difference between the spike and the distribution mean of the matched spikes was used as a measure for how different the propagation detected on this loose electrode was compared to the agreed upon spikes by RT-Sort and the other sorter, with absolute differences clipped at a maximum of 7 frames (Fig. 4C). Next, an interelectrode interval difference score per unit was defined as the average absolute difference over all spikes on all loose electrodes. To account for variability intrinsic to the variable action potential propagation speed, the average absolute difference over all matched spikes was subtracted from the average absolute difference of both the “RT-Sort only” and the “Other sorter only” spikes to get the final interelectrode interval difference score for both groups.

A paired one-sided t-test showed that the interelectrode interval difference score was significantly larger for each of the other sorters compared to the RT-Sort only spikes, taken over all units with an overlap score of ≥0.1 where both sorters detected at least 10 unique spikes (Fig. 4DE, SpyKing Circus: P=3.50*10^-7^, n=115 units; Kilosort2: P=1.49*10^-8^, n=119 units; Herdingspikes2: P=9.76*10^-5^, n=78 units; Tridesclous: P=5.87*10^-10^, n=104 units; HD-Sort: P=2.36*10^-13^, n=108 units; IronClust: P=2.43*10^-4^, n=106 units). Note that n is notably lower than the number of units detected by these sorters which is due to the fact that RT-Sort has a lower recall on unit detection due to more stringent detection thresholds (Sup. Fig. 8A).

We applied the same procedure on detected spike amplitude differences instead of interelectrode interval differences. Also here, we noted a significantly larger difference score for the “Other sorter only” spikes compared to the “RT-Sort only” spikes, except for HDSort (Fig. 4FG, SpyKing Circus: P=1.15*10^-8^, Kilosort2: P=2.49*10^-8^, Herdingspikes2: P=5.83*10^-4^, Tridesclous: P=5.55*10^-7^, HD-Sort: P=0.0789, IronClust: P=8.24*10^-6^). However, HDSort had the largest increase in the interelectrode interval difference score compared to RT-Sort of all other sorters.

Despite there not being a ground truth for these recordings, these results indicate that the RT-Sort only detections were more similar to the matched detections than the detections that were only made by the other sorter. This makes it more likely that the RT-Sort only detections reflect false negatives of the other sorter while the other sorter only detections could reflect false positives of that sorter. And so we conclude that RT-Sort is able to detect a lower number of units but that the units that are detected are of relatively high quality. Most importantly, these results were obtained by detecting spikes in online mode where it took 7.62ms±1.58ms (mean±STD) between the trough of the waveform and its sorted detection (Fig. 4H).

Altogether, our results indicate that RT-Sort performs at least at a comparable level as the currently available spike sorting algorithms, although with a lower unit recall. However, unlike all of the other spike sorting algorithms, RT-Sort is able to achieve this performance in real time, with sorting occurring within milliseconds after the spike occurred, while the experiment is still ongoing. Thus it is the first algorithm that allows for closed loop feedback experiments with single neuron resolution and mono-synaptic delay times.

## Discussion

The technological advancements in hardware used for extracellular neural recordings have yielded dense multi electrode recording devices for both *in vivo* (Jun et al. 2017, Ye et al. 2023) and *in vitro* (Ballini et al. 2014) recordings with interelectrode pitches smaller than 20µm. These devices allow for the recording of the activity of a single neuron by multiple adjacent electrodes.

With the high temporal resolution of these recording devices in the range of 20-30 thousand frames per second, consistent time delays can be observed among electrodes that detect the action potential of a single neuron resulting from the time it takes for this action potential to propagate along the axon (Tovar et al. 2018). This action potential propagation detection has been used to enhance the classification of neuron types based on high density recordings (Jia et al. 2019). In this work, we presented RT-Sort (Real Time Sorting) that for the first time utilizes action potential propagation detection instead of spike waveform template matching as a spike sorting methodology to identify individual neurons in high-density extracellular field recordings.

Spike sorting based on action potential propagation detection overcomes the issues related to changes in waveform shapes and overlapping spikes (Fig. 2, Sup. Fig. 4, 5). Simultaneously, action potential propagation based spike sorting reduces the rate of false positive spike detections (Fig. 2&4). Importantly, all of this can be achieved while being able to detect and sort spikes within less than 10ms after their trough time (Fig. 3), as opposed to having to wait until completing a recording before the spike sorting procedure can even be initiated. This puts the sorting latencies of RT-Sort in the range of signal transmission delay times of single synapses (Koch et al. 1996).

To obtain an optimal sorting performance, it is important to detect the propagating action potential on as many electrodes as possible, with high temporal precision. Therefore, we developed a convolutional neural network (CNN) that detects waveform shapes in noisy traces on single electrodes (Fig 1A-D, Sup. Fig. 1A-D), with a significantly higher recall than 5RMS based spike detection (Fig 1G, Sup. Fig. 1G). When training our model, we ensured that it is sensitive to the wide variety of waveform shapes that can occur while the action potential propagates along various parts of a neuron. As a result, even the *in vivo* recordings from Neuropixels probes yielded propagating action potentials over 150µm, taking up to 500µs for their propagation (Sup. Fig. 8BC).

The single electrode CNN detections form the inputs for the subsequent spike sorting algorithm. In offline mode, this algorithm takes the spike detections from a recording and finds recurring sequential activations of adjacent electrodes in sub-millisecond time scales. Using the interelectrode intervals of these observed sequential detections, newly occurring action potentials in live ongoing recordings can directly be matched to the previously detected propagation pathways, resulting in real time spike sorting, which is the basis for designating the algorithm RT-Sort (Real Time Sorting). The only requirement for applying RT-Sort in online mode is to make a short pe-recording of up to 5 minutes, using the same electrode configuration as will be used in the actual experiment. Subsequently processing this recording in offline mode to detect the sequences that can be used in online mode takes about 1.93±0.30 minutes (mean±STD) per minute of recording (Fig. 3G).

In this work, we presented a version of RT-Sort optimized for organoid recordings from HD-MEAs from Maxwell Biosystems (Fig. 1&2) (Ballini et al. 2014) and a separate version of RT-Sort optimized for mouse recordings from Neuropixels 1.0 (Sup. Fig. 1, Fig. 4) (Jun et al. 2017). However, these are just examples of commonly used recording devices for both *in vitro* and *in vivo* applications. RT-Sort was developed to easily produce optimized versions for any dense multi electrode recording device with interelectrode spacing of ≤50µm. The main aspect of recording device and biological sample specific optimizations for RT-Sort resides in the single electrode spike detection model. Here we trained a CNN to optimally detect waveforms from a specific biological model type in recording device specific noise (See *Methods* for more details). If another biological sample type and/or recording device were to be used, the currently presented versions of RT-Sort might still yield good performance. For example, we successfully applied our human organoid MEA model on rat primary cultures plated on these same MEAs (Fig. 3A-H), and we got good results for a subset of units when we applied our mouse Neuropixels model on biophysically simulated waveforms in random noise (Fig. 3I-J). However, for optimal results, we recommend training a model that is specific to the biological sample type and recording device. This can easily be done through a Python library that will soon be released publicly and only requires a few previous recordings from the same biological sample type and recording device combination.

RT-Sort has been tested on a simulated ground truth recording as well as simultaneous intracellular and extracellular field ground truth recordings. Furthermore, its detection and sorting performance has been compared on real *in vivo* recordings (Siegle et al. 2021) against a variety of commonly used spike sorters (Hilgen et al. 2017, Pachitariu et al. 2018, Jun et al. 2017b, Yger et al. 2018, Garcia & Pouzat 2015, Diggelmann et al. 2018). Our results indicate that RT-Sort has a high precision, meaning that it detects very few false positive units and spikes (Fig. 2). In the recording without ground truth, this is reflected in the fact that the spikes that RT-Sort detects but that are not detected by the other sorter are more similar to the spikes that both sorters agree on compared to the spikes that the other sorter detects but that RT-Sort doesn’t (Fig. 4A-G). This makes it more likely that the RT-Sort only detections reflect false negatives of the other sorter while the other sorter only detections could reflect false positives of the other sorter. These results come with the price of a lower recall on unit detection, meaning that RT-Sort misses units that other sorters might detect. Since our recall of single spike detections was significantly higher than 5RMS (Fig. 1G, Sup. Fig. 1G), we suspect that this reduced recall on unit detection is due to the more stringent unit detection bounds imposed on RT-Sort, requiring the detection of a spike on at least 4 different recording electrodes. These bounds are applied to force detections to resemble propagating action potentials which leads to the improved precision.

Given that the main differentiator of RT-Sort is its short latency spike sorting in the millisecond range, we anticipate that the lower unit recall does not form a significant issue for the experimenter as long as the detected units are sorted with the high accuracy that we report. This would still enable the experimenter to design a wide range of closed loop experiments with tens to hundreds of units that can be recorded in real time (RT-Sort detected 168 units in the Neuropixels recording in Fig. 4). Nonetheless, we anticipate that future iterations of action potential propagation-based spike sorting algorithms will further improve on RT-Sort’s performance, both when it comes to precision and recall as well as the sorting speed.

Areas where we see further space for improvement include the architecture and training procedure of the single spike detection model. For example, we have observed slightly better performance for deeper detection models (consisting of more layers) but this also notably increased detection latencies. A better detection performance might also be obtained by including a larger number of averaged waveform shapes used for generating semi-artificial training data to get an even more complete picture of all the different possible waveform shapes that might be present in a recording. Furthermore, there will likely be room for further optimizations in the sorting algorithm and chosen parameters for detecting propagation sequences, merging them and assigning spikes. For example, to better handle electrode drift in long duration recordings. Finally, through clever multithreading and enhanced computational processing power, we anticipate the sorting latencies of future propagation-based spike sorting routines to approach latencies below 5ms.

In making improvements, one will have to carefully balance sorting accuracy with detection latencies. With the continuous improvement of processing capabilities, we expect this balance to increasingly favor sorting accuracy without having to give in on detection latencies. Note that all the reported results were obtained using a single NVIDIA-RTX-A5000 card, which we anticipate to soon become outdated and easy to outperform. In presenting RT-Sort, we hope to evoke a paradigm shift in the field of spike sorting towards propagation based methods over template matching methods.

Despite leaving space for improvements, we think that in its current form, RT-Sort will already provide significant value to the researchers that conduct neural extracellular field recordings with dense multi electrode recording devices, both *in vivo* and *in vitro*. RT-Sort enables, for the first time, an easily implementable spike sorting routine with time latencies for sorted spike detection that are comparable to single synaptic delay times. This important feature of RT-Sort opens up interesting opportunities for closed loop experiments, where a neural system receives external inputs based on preceding activity recorded from the same system. These types of experiments have been performed in various forms but always utilize multi-unit activity as read-out of the neural system (DeMarse et al. 2001, Wagenaar et al. 2005, Bakkum et al. 2007, Kagan et al. 2022). With RT-Sort, closed loop experiments can now be conducted with real time single unit read out accuracy, which enables studying neural circuit dynamics in response to external inputs with a greatly enhanced granularity. Similarly, we anticipate that this single unit real time detection granularity for large populations of recorded neurons can greatly enhance the performance of brain machine interfaces utilized in real time.

In summary, RT-Sort exemplifies a novel approach to spike sorting based on action potential propagation which comprises high-fidelity sequential detection of a propagating action potential on multiple adjacent electrodes with sub millisecond but nonzero time delays. This approach lies closer to the biology that underlies action potentials than template matching, the current norm in spike sorting algorithms. As a result, RT-Sort is more robust against waveform shape changes and overlapping waveforms than conventional methods, yielding a reduced false positive rate in detected units and spikes. Importantly, RT-Sort is able to detect and sort spikes within less than 10ms after their occurrence, on the range of single synaptic delay times. This makes it an interesting tool for closed loop experiments and brain machine interfaces as well as various other implementations. We hope that the broader community will take interest in action potential propagation-based spike sorting and apply this new paradigm to continuously iterate on better performing real time spike detection algorithms.

## Methods

### Real time spike detection

RT-Sort uses action potential propagation detection to identify individual neurons in recordings with dense electrode configurations of ≤50µm interelectrode pitch. This requires a sensitive detection of the action potential on multiple electrodes followed by a sorting procedure that assigns the action potentials to the right neurons. All of this needs to happen in millisecond timescales to be able to use RT-Sort in real time. For this, we developed a spike detector that can simultaneously detect spikes on over a thousand different electrodes within millisecond time scales, while being more sensitive than 5RMS threshold crossings and able to detect overlapping waveforms.

A convolutional neural network (CNN) was trained to detect neural waveform shapes in a noisy signal. Due to variability in the type of noise produced by different recording devices, it is important to train a specific model per recording device type. In this work, separate models are trained for human brain organoids recorded with Maxwell Biosystems multi-electrode arrays (Ballini et al. 2014) and for intact mouse brains recorded with Neuropixels probes (Jun et al. 2017). The data used for the training and validating process comes from (Sharf et al. 2022) and (Siegle et al. 2021) respectively and can be found here https://datadryad.org/stash/dataset/doi:10.25349/D9031Z and here https://buzsakilab.nyumc.org/datasets/SiegleJ/AllenInstitute_744912849/session_766640955/. All training and validating is done using an NVIDIA-RTX-A5000 GPU.

### CNN architecture

The CNN for spike detection consists of 4 convolutional layers. The first three convolution layers have 50 filters with a kernel size of the number of frames in 1 millisecond of raw data + 1 (21 for MEAs with sampling rate of 20kHz, 31 for Neuropixels with sampling rate of 30kHz), stride of 1, and ReLU nonlinearity. The last layer (the output layer) has 1 filter with the same kernel size and stride as the previous layers and followed by a Sigmoid nonlinearity (Fig 1A, Sup. Fig. 1A for Neuropixels model with slightly altered parameters to account for different sampling rate). The CNN takes in a 10ms snippet of raw, unfiltered data. The output layer contains a node for each frame in the 2-8ms period within the input data. The kernel size was chosen so that each node in the output layer receives information from all frames in the input layer in the period of 2ms before and after the frame that the node makes the prediction for.

With this architecture, the model is trained so that for the provided snippet of 10ms of data, it returns a likelihood of a spike waveform trough to be present for each recording frame in the period of 2-8ms within this window. The outer boundaries of the window are not considered since the full shape of the waveform would not be present within the window, leading to a worse detection performance. Each frame receives a score between 0 and 100 indicating the likelihood of there being a waveform trough in this frame. Before each 10ms sample is fed into the CNN for training or inference, it is converted from int16 format to float16 and standardized by subtracting the median of the sample.

### Spike detection semi-artificial data generation

Note that it is difficult to obtain sufficient ground truth spike detections to effectively train a neural network. The availability of high electrode density electrophysiology recordings with corresponding ground truth is limited, and it is difficult to generate this data (Steinmetz et al. 2018). Alternatively, ground truth training and validation datasets are generated through artificial data amplification, inspired by the work from Parks et al. (Parks et al. 2018). First, a selection of recordings from the recording device and sample type of interest are made. In our case, 6 recordings from different human brain organoid slices are selected where 5 recordings are used to generate training data, and 1 recording is used to generate validation data in a 6-fold cross validation manner as described next. A same selection is made for intact mouse brain recordings made *in vivo*. Note that during validation, only data generated from a different replica of the recording device is used (although from the same type), interfacing with a different biological sample (organoid slice or mouse brain). In this way, the model is trained for recording device type and sample type specificity while allowing for generalizability among different replicates of recording devices of the same type and generalizability among different biological replicates.

The selected recordings are bandpass filtered between 300Hz and 6000Hz and subsequently spike sorted using Kilosort2 and curated. In the initial curation round, all units with at least 30 detected spikes, less than 1% of spikes violating a <1.5ms inter spike interval and a signal to noise ratio larger than 5 on the main electrode are selected. The spikes of these units are considered to be all spikes in the recording. From the selected units, a smaller selection is made for training our spike detection model by applying a second round of curation. Here all units with at least 50 detected spikes are selected and the filtered signal on each electrode is averaged around the maximum absolute waveform peak from the detections on the main electrode.

In order to detect a propagating action potential along the whole axon, it is important for the spike detection algorithm to be able to detect spikes that are not just from the highest amplitude electrode. Therefore, waveforms were selected from all electrodes where the detection trough of the averaged waveforms cross −19µV (−36µV for the Neuropixels data). Alternatively, if the electrode detected a waveform with a peak at least twice as large as the through, this electrode was selected if the peak crossed 19µV (36µV for the Neuropixels data). Also, the standard deviation in the peak amplitude divided by the absolute value of the amplitude had to be less than or equal to 0.6. The averaged waveforms for these electrodes are stored for generating training and testing samples (Fig. 1B, Sup. Fig. 1B for Neuropixels example). This yields 82.83±74.62 averaged waveforms from single electrodes per recording (1327.33±427.00 for the Neuropixels). Note that the selection includes averaged waveforms of a broad variety of shapes from electrodes that recorded a signal from any part of a neuron, which allows us to detect a propagating action potential along different parts of the neuron.

From the same recordings as used to select the average waveforms, 10ms periods with no detected spikes on any electrode are extracted to obtain recording device specific noise. To ensure that there is no presence of waveforms in these 10ms periods, only periods where there were no spike troughs detected in the 3ms before the start of the window until 3ms until after the end of the window were used. Spikes after the initial curation round were used to find these no-activity periods. Next, an averaged waveform is pasted into a noise snippet to create an artificial spike event that can be used for training and validation (Fig. 1C, Sup. Fig. 1C for Neuropixels example). In this way, the average waveform shapes pasted into the recording and background noise allowed us to generate near infinite training samples that we could use to train our CNN to be able to detect waveform shapes of any part of a neuron in noise intrinsic to the sample preparation and recording device. Furthermore, multiple waveforms can be pasted into the same noise snippet to artificially create overlapping waveforms. When doing so, the number of pasted waveforms was randomly determined for each generated sample, following the probability distribution 50%, 30%, 12%, 6%, 2% for 0, 1, 2, 3, or 4 waveforms respectively. The distance between waveform troughs was randomly determined within a range of 0.2ms to 6ms apart (Fig 1D, Sup. Fig. 1D for Neuropixels example). In cases where multiple waveforms are added to the same noise snippet, no sample has multiple copies of the same waveform shape.

During the 6-fold cross-validation training, one of the 6 recordings is used as the validation dataset while the remaining 5 are used as the training dataset. For the MEA model, one validation and training epoch consisted of 2 and 20 samples per waveform in the dataset, respectively. For the Neuropixels, only 1 sample per waveform was used since recordings yielded more different waveform shapes for the training and validation datasets. In the training dataset, a waveform from any of the 5 recordings could be pasted into a 10ms noise sample from any of the other 5 recordings.

### CNN training

The generated training samples are used to train the aforementioned CNN. The batch size is 1. Stochastic gradient descent with an initial learning rate of 0.000776 (0.0002 for the Neuropixels model) and momentum factor of 0.85 is used. If the validation does not decrease by at least 0.0001 (0.01 for the Neuropixels model) after 5 epochs, the learning rate decreases by a factor of 0.4. These optimal hyperparameters were obtained through 6-fold cross-validation with Bayesian tuning using CometML. The model is trained until the validation loss does not decrease by at least 0.01 after 10 epochs. After training is completed, the weights that give the best validation loss are used. This yields 33.5±7.1 epochs (mean±STD) for the MEA model and 18.7±5.0 epochs for the Neuropixels model. The small differences between the training loss and testing loss indicated that our models are not overfitting nor underfitting the data (Fig. 1E, Sup. Fig. 1E for Neuropixels results).

### Validating and detection threshold

Model detections within 0.4ms of the actual absolute waveform peaks from the samples pasted into the snippet are considered to be true positives. For a range of thresholds for the detection score, the precision and recall are computed as a function of the detection threshold. The optimal detection threshold of each model trained on one of the 6 folds was chosen by optimizing the F1 score. Using this threshold, the performance of the model detections compared to 5RMS based detection was assessed by computing the precision and recall.

The training and validation were performed using semi-artificially generated samples as described above. The MEA model trained on the fold with the largest number of waveforms in the training dataset and the Neuropixels model trained on the fold where the validation dataset consisted of the recording described in *Results: RT-Sort compared to other sorters* were then applied in our real time spike sorting algorithm as described below. The results after sorting were further tested against actual ground truth datasets as will be described in *Methods: Real time spike sorting*.

Since for the spike sorting algorithm, we require sequential detections on a specific set of electrodes in a specific order, detecting false positives on single electrodes is less problematic than detecting false negatives. A false positive detection on a single electrode will not yield an actual detected spike since it will not meet the sorting thresholds. As a result, we will apply two model detection thresholds: the stringent threshold (which is at the optimal F1 score) and the loose threshold (below the optimal F1 score). Specifically, we will use a stringent threshold of 27.5% (17.5% for the Neuropixels model) and a loose threshold of 10% (7.5% for the Neuropixels model). The loose threshold was chosen such that the recall on the validation dataset was about 15% larger than the precision, in order to prioritize a low false negative rate of single electrode spike detection. These thresholds can be adjusted by the user based on their sample and their experimental needs.

### CNN inference

The trained model is applied to recordings with varying degrees of noise. To ensure that the model’s inputs are on the same amplitude scales as it encountered during training, the mean amplitude deviation of the 5 recordings in its training dataset (training amplitude deviation) is calculated. The amplitude deviation of a recording is calculated as follows: for the first 50ms on each electrode, the interquartile range of the recording traces is calculated. The median of these ranges is the amplitude deviation.

To scale a recording to the model’s expected amplitude range, the amplitude deviation of the recording is calculated, and the recording’s traces are multiplied by the training amplitude deviation divided by the recording’s training amplitude deviation.

### Real time spike sorting core parameters

The sorting algorithm takes the raw recording traces and real time detection scores as inputs and finds propagating action potentials based on recurring consecutive detection patterns on adjacent electrodes. The algorithm can be used to process previously made recordings (offline mode), and it can also be applied in real time (online mode). Using RT-Sort for real time sorting requires at least a 40-second pre-recording (5 minutes is ideal for optimal sequence detection) with the same electrode configuration as the actual experiment to be made before the experiment in order to detect sequences. Afterwards, action potentials detected in real time are assigned to the sequences detected in the pre-recording.

For the description of the real time spike sorting algorithm below, the stringent and loose detection thresholds will be used as defined previously. Furthermore, two types of distance thresholds will be used. The “inner” electrodes will be within 50µm of the electrode of interest whereas the “outer” electrodes will be within 100µm of the electrode of interest. These distance metrics were chosen for optimal performance with a checkerboard Neuropixels configuration as well as Maxwell MEA electrode configurations that use every other electrode resulting from a sparse 7x activity scan, yielding a 35µm pitch (Sup. Fig. 9). Action potential propagations are first detected based on the pre-recording and then used for real time spike detection and sorting in online mode in the actual experiment.

### Propagation sequence detection

The first phase of the sorting algorithm consists of propagation sequence detection. For each recorded electrode, which will be referred to as the root electrode of the sequence (Fig 2A, marked with red star), all stringent model detections are obtained. Only the stringent model detections that have their largest standardized amplitude (defined below) on the root electrode and no higher standardized amplitude detections within 0.5ms on any of its outer electrodes are considered.

All amplitudes are detected in the raw, unfiltered data. To account for LFP impacting the amplitude, amplitudes are standardized according to the calculation below. In the remainder of the text, any reference to amplitudes pertains to amplitudes standardized in this manner:

The voltage at the model’s predicted spike location (“absolute amplitude") on the electrode is extracted. For the preceding 50ms (“window”), the following values are found:

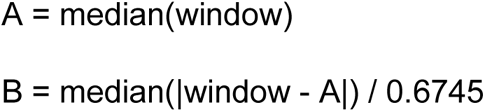

The standardized amplitude is then equal to (absolute amplitude - A) / max(0.5, B). Where B is an estimate of the standard deviation of the background noise (Donoho & Johnstone 1994, Quiroga et al. 2004).

The selected model detections on the root electrode that have a stringent detection on at least one inner electrode within 0.1ms are selected. These will be referred to as the root detections. All root electrodes with an average firing rate of at least 0.05Hz over the whole recording are selected for further processing. For each root electrode, all root detections are compared to all stringent detections on the inner electrodes of the root electrode in a pairwise manner. In this comparison, the root detections might be split into multiple clusters as follows:

During the comparison to each inner electrode (“the splitting electrode”) the time intervals between the root detections and the detections on the splitting electrode that occur within 0.5ms of any of the root detections (“splitting codetections”) are obtained. 1-component, 2-component, 3-component, and 4-component Gaussian mixture models are fit to the codetection interval distribution. The model with the lowest Bayesian information criterion is used to determine whether there are a single or multiple interelectrode propagations present in the interval distribution (Sup. Fig. 10A). To prevent overfitting the distribution, a Gaussian mixture model is ignored if it has a negative Bayesian information criterion (BIC). To prevent underfitting the distribution, a Gaussian mixture is ignored if the fitting of the model yields a FloatingPointError due to encountered underflow. If all Gaussian mixtures are ignored, the 1-component Gaussian mixture model is used.

Finally, any cluster that has fewer detections than max(10, 10% of the root detections) are discarded and their splitting codetections can be used to form other clusters (see below). If all clusters do not meet this threshold, then the root detections remain together as a group and are not split into clusters.

The root electrode’s inner electrodes are sorted in ascending order of distance. Let the i^th^ electrode refer to the i^th^ electrode in this sorted order. All root detections are first compared to the i^th^ electrode. Every cluster that is formed is then compared to the i+1^th^ electrode, treating each cluster the same as a set of root detections as described previously. This process repeats until no more electrodes remain, resulting in a branching process of the root detections splitting into clusters which then split into more clusters based on differences in interelectrode intervals. Each root detection can only join one cluster. After the clusters starting on the splitting with the i^th^ electrode have been fully formed, the remaining root detections not assigned to a cluster are then compared to the i+1^th^ electrode. This process repeats until there are no more root detections or no more inner electrodes for direct comparison with the root electrode.

To accommodate long neurons that could extend farther than the root electrode’s inner electrodes, the list of splitting electrodes for splitting a cluster can expand. If at least 50% of the root detections form a single cluster, then the inner electrodes of the most recent splitting electrodes are added to the list of splitting electrodes in ascending order of distance with the most recent splitting electrode. To prevent a single cluster from traversing the entire array, a patience counter is used. The patience counter starts at 0. If a cluster is not split (i.e. the root times stay together) after all the splitting electrodes with the same distance are considered, then the patience counter increases by 1. If it reaches 6, there are no more attempts to split the cluster, regardless of the potentially remaining splitting electrodes. If the cluster is split, the patience counter resets to 0. The clusters formed in the previous steps are each considered to be preliminary propagation sequences.

It is possible for one preliminary propagation sequence to contain the spikes from multiple neurons if the spikes have very similar intervals on multiple electrodes. To account for this, a distribution is constructed of the amplitudes on the root electrode. If a Hartigan’s dip test indicates that this amplitude distribution is not unimodal (P<0.1), the preliminary propagation sequence is split into multiple sequences. To do this, 2-component, 3-component, and 4-component Gaussian mixture models are fit to the amplitude distribution (using the GaussianMixture object from the Python scikit-learn library). Only the model with the lowest BIC is used to split the amplitude distribution (Sup. Fig. 10B). See Sup. Fig. 10C for the distribution of all P-values resulting from applying the Hartigan’s dip test on the root electrode amplitude distributions. After splitting, sequences with less than 10 spikes are discarded.

### Extracting the detections, intervals, and amplitudes

Next, the median of the detection scores for all detections in a preliminary propagation sequence are computed for each electrode that is part of the preliminary propagation sequence or that falls within 50µm of any electrode in the preliminary propagation sequence. Based on the median detection scores, each sequence will have three groups of electrodes:

1. Loose electrodes: electrodes that have a median detection above the loose threshold and that are within 50µm of any of the preliminary propagation sequence’s splitting electrodes (including the root electrode)
2. Inner loose electrodes: loose electrodes that are within 50µm of the root electrode
3. Footprint electrodes: loose electrodes and any electrode within 50µm of a loose electrode

Any sequence that has fewer than 4 loose electrodes or fewer than 3 inner loose electrodes is discarded to meet spike assignment thresholds (see *Methods: Spike reassignment*).

For all footprint electrodes, 0.5ms of the signal before and after the root electrode detection time point is extracted for each of the action potential propagation sequence detections in the preliminary propagation sequence (“propagation period”). The model’s detection outputs are obtained for this propagation period. Iterating over all detections in the preliminary propagation, the maximum value in the detection output during the propagation period is obtained for each footprint electrode. The time delay between this maximum detection and the maximum detection time on the root electrode is considered to be the electrode’s interval for that detection. The amplitude at the electrode’s maximum detection is considered to be the electrode’s amplitude for that detection. The median detection score, mean interval, and mean amplitude across all detections are found individually for each footprint electrode (Fig 2A, waveform colors reflect latencies). If an electrode’s median detection score is less than 3%, it is set to 0. Finally, the standard deviation in the root electrode’s amplitudes of all detections are extracted.

To remove sequences that are the result of noise spikes across all electrodes, any sequence that has more than max(100, 10% of the sampled electrodes in the recording) electrodes with a median detection score above the loose threshold is discarded.

### Spike reassignment

The preliminary propagation sequences might only contain a low number of detections due to the strict detection thresholds. Therefore, spikes are reassigned to the preliminary propagation sequences using the same pre-recording that was used to detect the preliminary propagation sequences, in order to make the extracted median detections, mean interelectrode intervals, and mean amplitudes more robust and representative of the actual neuron that underlies the preliminary propagation sequence.

A spike may be assigned to a preliminary propagation sequence if all the following conditions are met:

1. A stringent detection must occur on the root electrode.
2. The amplitude of this detection on the root electrode must fall within 2.5 standard deviations of the mean amplitude on the root electrode.
3. At least 3 of the inner loose electrodes and max(4, 33% of the number of loose electrodes for the sequence) of the loose electrodes need to detect a spike above the loose threshold within the propagation period.
4. For all footprint electrodes, both the weighted interelectrode interval difference (relative to the root electrode) and the weighted amplitude percent difference (relative to the sequence’s mean amplitudes) cannot be greater than the interelectrode interval and amplitude thresholds, respectively. For MEAs, the thresholds are 3.5 frames and 65% respectively (Fig. 2D-F for 3 single spike detections). For Neuropixels, the thresholds are 2.5 frames and 45%, respectively (Sup. Fig. 4-6 for examples of single spike detections).

The “weight” of each footprint electrode is calculated as its median detection score divided by the sum of all the footprint electrode’s median detection scores. Subsequently, for each footprint electrode, the difference (or percent difference) between the sequence’s mean interelectrode interval (or amplitude) and the spike’s interval (or amplitude) on that electrode is found. Differences are clipped to be at most twice the interelectrode interval and amplitude thresholds. Then, the weighted sum of the differences is calculated using each electrode’s weight.

When multiple sequences with the same root electrode detect a spike at the same time, the spike’s difference score with each sequence is found. The difference score is calculated as the sum of:

1. The interelectrode interval difference divided by the interval threshold
2. The amplitude difference divided by the amplitude threshold
3. -0.5 times the number of the sequence’s loose electrodes that detect the spike divided by the number of electrodes that have a loose threshold crossing

The spike is only assigned to the sequence with the lowest score. This process will be referred to as “repeated detection removal”. After reassigning the spikes, new median detection scores, mean interelectrode intervals and mean amplitudes are computed for each preliminary propagation sequence. Preliminary propagation sequences that, after spike reassignment, have less than 10 spikes or a firing rate less than 0.05Hz are discarded.

### Merging

Detecting the preliminary propagation sequences in this manner allows for the possibility of detecting the same axon multiple times. For example, the first electrode in the preliminary propagation sequence that detects a signal after the root electrode might also be detected as a separate root electrode for the same axon. Therefore, the next phase of the algorithm consists of merging the detected preliminary propagation sequences into the final propagation sequences. For the first merging round, only preliminary propagation sequences that share the same root electrode are considered. For all footprint electrodes, the interelectrode intervals and amplitudes compared to the root electrode, weighted by their average detection score, are compared. The following conditions must be met for two sequences to be merged into one:

1. At least 3 electrodes must be in both sequence’s inner loose electrodes.
2. The number of electrodes in both sequence’s loose electrodes must be greater than max(4, 33% of the total number of electrodes that are loose electrodes for at least one of the sequences).
3. The weighted interelectrode interval difference for both sequence’s footprint electrodes is at most 3.5 frames for MEAs and 2.5 frames for Neuropixels. The same clipping as in spike reassignment is used.
4. The weighted amplitude percent difference for both sequence’s footprint electrodes is at most 65% for MEAs and 45% for Neuropixels. The same clipping as in spike reassignment is used.
5. The absolute difference between the sequences’ mean amplitudes on the root electrode is calculated, and the largest standard deviation in the amplitudes between the sequences is found. The difference divided by this standard deviation must be at most 2.5. Due to higher neuron density in the *in vivo* recordings, this condition is implemented for Neuropixels only. The user can choose whether to use this requirement or not.

If the two sequences are merged into one, the median detections, mean interelectrode intervals, and mean amplitudes are updated accordingly.

The weights and differences for the root electrode are not considered for the weighted interelectrode interval difference since the intervals are relative to the root electrode, making the interval always 0. The midpoint method is used for the amplitude percent difference. The merge score is the sum of the interelectrode interval difference divided by the interval threshold and the amplitude percent difference divided by the amplitude threshold. The pair of preliminary propagation sequences with the lowest score is merged first until no more pairs can be merged or there is only one sequence remaining.

To better asses sequences for merging in the subsequent merging round, the root electrode of each sequence is replaced by an electrode with a mean amplitude of at least 80% of the current root electrode’s mean amplitude, a median detection score above the stringent threshold, and a interelectrode interval less than −2 frames. If multiple electrodes meet these criteria, the electrode with the most negative interelectrode interval becomes the root electrode. If no electrodes meet these criteria, the current root electrode remains as the root electrode. The interelectrode intervals on other electrodes are updated accordingly.

In the subsequent merging round, root electrodes of a preliminary propagation sequence that are inner electrodes of a root electrode from a different preliminary propagation sequence are assessed for merging in the same way as described above. In this case, the root electrode with the highest mean amplitude is used as a reference for the interelectrode interval difference, and for Neuropixels, the root electrodes of both sequences must pass condition #5 described above. This final round of merging yields the actual propagation sequences. For each of these sequences, the electrode with the highest median detection score becomes the root electrode, and the intervals on the other electrodes are updated accordingly.

To conclude the pre-experiment preparations, spikes are reassigned to sequences in the same way as described in *Methods: Spike reassignment* but this time with applying repeated detection removal to all groups of root electrodes that detect a spike within 0.2ms and that are inner electrodes to each other. In the case of using RT-Sort for offline spike sorting, these spike times are the final results.

### Real time spike sorting

In the case of using RT-Sort on online mode for real time spike sorting, the same process as spike reassignment is used to spike sort in real time using the updated median detection scores, mean interelectrode interval and mean amplitudes. Also here, repeated detection removal is applied to all groups of root electrodes that detect a spike within 0.2ms and that are inner electrodes to each other. Using the streamed data, a 10ms sample is processed every 5ms to account for the spikes in the first and last 2.5ms parts of the sample that can’t be detected due to biological constraints (Fig. 3A).

### Patch-MEA ground truth recordings

To test the performance of RT-Sort, a ground truth dataset was generated by recording from a primary rat culture plated on a Maxwell MEA while simultaneously, a patch clamp recording was obtained from a single neuron. Spiking activity of the patched neuron was recorded in cell-attached mode or whole-cell current clamp and was also detected by the MEA. The patch-MEA dataset from (Bartram et al. 2023) was used, as well as an additional dataset, generated largely according to the same protocol using only the whole-cell current-clamp mode and internal solution for recording. The experimental protocols involving animal tissue harvesting were approved by the veterinary office of the Canton Basel-Stadt according to Swiss federal laws on animal welfare and were carried out in accordance with the approved guidelines. Cell culturing protocols are identical to those described in (Bartram et al. 2023). Both datasets contained 2 individually patched neurons. One of the two datasets contained an additional 5-minute pre-recording that was made using the MEA only, which allowed for the detection of propagation sequences in order for RT-Sort to be run in online mode on the patch-MEA recording. This pre-recording occurred within 60 minutes before the simultaneous patch-MEA recordings.

The simultaneous patch-MEA recordings have durations ranging from 40 seconds to over 3 minutes. By replaying these recordings, a real time application of RT-Sort was mimicked. Intracellular voltage peaks above 4x RMS of the patch recordings are considered to be spike events. RT-Sort spike detections are compared to these patch spike events by computing an overlap score for each of the 4 patched neurons.

### Overlap score

Following (Buccino et al. 2020), when two spike trains (A and B) are compared to see if they detect a signal from the same neuron, an overlap score is computed as follows:

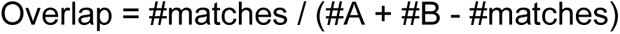

Here #matches are the total number of spikes from the two spike trains that are detected within 0.4ms of each other, #A are the total number of detected spikes in spike train A and #B are the total number of detected spikes in spike train B. In the case of the patch-MEA ground truth recordings as described in *Methods: Patch-MEA ground truth recordings*, the patch spikes are considered to be spike train A while the RT-Sort spikes are considered to be spike train B. In the case of the sorter comparison on the Neuropixels recording as described in *Methods: Sorter comparison on in vivo Neuropixels recording*, the overlap score is used for all pairwise comparisons between sorters.

### Simulated Neuropixels ground truth recording

Genuine ground truth recordings obtained by combining extracellular field recordings and patch clamp recordings are hard to obtain *in vitro*, and even harder *in vivo* (Steinmetz et al. 2018). This yields a low number of ground truth units while these units are often biased to be larger in size and amplitude and higher in their firing rates. As a result, these units are relatively easy to sort and might not form a representative sample of all neurons that are part of a large-scale recording.

As a result, we also tested RT-Sort on a simulated Neuropixels recording. This simulated ground truth dataset consisted of 250 biophysically detailed neurons that each exhibit independent Poisson firing of spikes with added Gaussian noise. See (Buccino et al. 2020) for a full description of the generation of the dataset. RT-Sort was first tested on this simulated recording using sequence metrics generated from the known ground-truth spikes times to focus the performance assessment on spike assignment to propagation sequences in online mode. Next, RT-Sort was used in offline mode on the first 5 minutes of the simulated recording to also detect the propagation sequences which were then used to test RT-Sort again in online mode on the second 5 minutes of the simulated recording.

### Sorter comparison on *in vivo* Neuropixels recording

In addition to a simulated Neuropixels recording, RT-Sort is tested on a genuine Neuropixels recording without ground truth. The recording was part of a larger dataset made from a head-fixed mouse by the Allen Institute for Brain Science (Siegle et al. 2021; Allen Institute for Brain Science, 2019; dataset ID: 766640955; probe ID: 77359232). We use the first 5 minutes of the recording to detect sequences as described in *Methods: Propagation sequence detection* and following sections. We then assign spikes to these detected sequences using the subsequent 5 minutes of the recording as described in *Methods: Real time spike sorting*. Other recordings from the same dataset are used to generate training data for our detection model as described in *Methods: Spike detection semi-artificial data generation*.

Neuropixels have a sampling frequency of 30kHz and can record from up to 384 channels simultaneously. This specific recording used a checkerboard configuration (Sup. Fig. 9) and yielded 246 active recording channels (the remainder of the channels were either not inserted in the brain tissue or had a firing rate below 0.1Hz). The probe records from part of the cortex (V1), the hippocampus (CA1), the dentate gyrus and the thalamus (LP). During the experiment, the mouse was presented with a variety of visual stimuli while freely running on a rotating disk (for more details see Siegle et al. 2021).

As shown in the work from Bucino et al. 2020, applying 6 different conventionally used spike sorters (HDSort, HerdingSpikes2, IronClust, Kilosort2, IronClust, SpyKING Circus and Tridesclous (Hilgen et al. 2017, Pachitariu et al. 2018, Jun et al. 2017b, Yger et al. 2018, Garcia & Pouzat 2015, Diggelmann et al. 2018)) to this recording yielded wildly varying results with limited overlap between the different sorters. However, without a ground truth, it is not possible to say which results are correct. We hypothesize that the low overlap between units could be due to false spike detections of different sorters which could make the spike trains of different sorters so different that they are not recognized as coming from the same unit. This would mean that good performance of a certain sorter could potentially be obscured by bad performance of a different sorter.

For that reason, in comparing RT-Sort to each of the 6 other spike sorters, we focused on the units detected by both RT-Sort and the other sorter (overlap score ≥0.1). For each of these units, a fraction of the spikes was detected by both RT-Sort and the other sorter while both RT-Sort and the other sorter also detected spikes that were not co-detected. For each unit that had at least 10 spikes in each of these 3 groups, the interelectrode intervals and amplitudes of the not co-detected spikes from each sorter were compared to the mean of the interelectrode interval and amplitude distributions over all co-detected spikes.

Specifically, for each of the loose electrodes except the root electrode, the interelectrode intervals relative to the root electrode and the amplitudes were obtained over every spike in each of the three groups. Next, the absolute difference in the interelectrode interval and the percent difference in the amplitude were computed for each of the spikes, in comparison to the distribution mean over all spikes that were detected by both sorters. The differences were then pooled over all loose electrodes which yielded three absolute difference distributions for the interelectrode intervals and three percent difference distributions for the amplitudes per unit. 1 distribution consisted of the differences for the co-detected spikes, 1 distribution consisted of the differences for the RT-Sort only spikes and 1 distribution consisted of the differences for the other sorter only spikes.

The mean of the RT-Sort only distribution and the mean of the other sorter only distribution were taken as a measure for how different these spikes were relative to the co-detected spikes. The mean of the distribution over all spikes that were detected by both sorters was taken as a measure for the intrinsic variability within all co-detected spikes. The mean over all co-detected spikes was then subtracted from both the mean over all RT-Sort only spikes and the mean over all other sorter only spikes to get a difference score for the RT-Sort only spikes and a difference score for the other sorter only spikes per unit. An individual score was computed for the interelectrode interval absolute differences and for the amplitude percent differences. Due to the normalization achieved by subtracting the mean of the differences of the co-detected spikes, a difference score of 0 indicates that the spikes in the distribution of either RT-Sort only spikes or other sorter only spikes were equally variable in their interelectrode intervals or amplitudes as the co-detected spikes. Meanwhile, difference scores larger than 0 indicate that the spikes in the distribution of either RT-Sort only spikes or other sorter only spikes were more variable in their interelectrode intervals or amplitudes as the co-detected spikes.

As a final comparison between RT-Sort and the other sorter, a paired t-test was performed to see whether either RT-Sort only spikes or other sorter only spikes were more different from the co-detected spikes, using the difference scores per unit. This t-test was computed separately for both interelectrode interval differences and amplitude differences.

## Author contributions

T.V.D.M. and K.S.K. designed, conceived and supervised the study with suggestions and comments from K.T., D.H. and D.F.P.; T.V.D.M., M.L. and D.C. developed the software and performed subsequent data analysis; J.B. and A.H. provided the patch-MEA ground truth recordings; A.R., L.R.P., P.K.H., D.F.P., J.B., K.T. and K.S.K provided feedback and ideas throughout the development process of RT-Sort; T.V.D.M. wrote the first version of the manuscript with support on the methods from M.L.; K.S.K. and P.K.H. provided valuable edits and suggestions on the subsequent drafts, and all authors discussed the results and commented on the manuscript.

## Acknowledgements

The authors would like to thank Alessio Buccino for sharing the results from applying the 6 different spike sorters to the Neuropixels recording used in Fig. 4 and for answering various of our questions throughout the development of RT-Sort. We would also like to thank Samantha Hermans for their work on related analyses and sharing their results.

