## Supplementary figures for "RT-Sort: an action potential propagation-based algorithm for real time spike detection and sorting with millisecond latencies"

This PDF file includes: Supplementary Figures 1 to 10

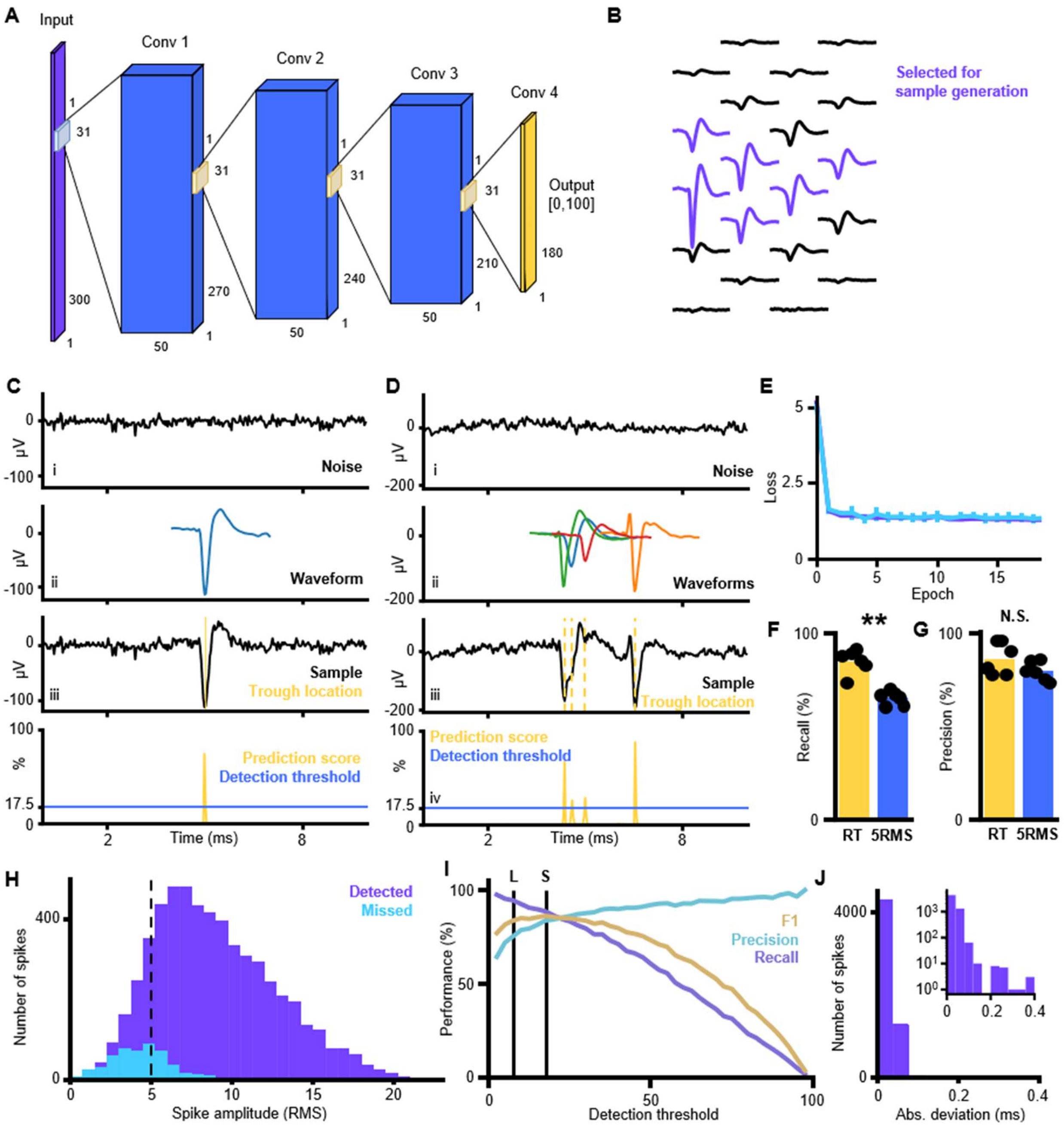


***Supplementary Figure 1: Action potential detection with high temporal accuracy using a convolutional neural network (CNN).*** *A) Architecture of the CNN used for spike detection in 30kHz Neuropixels recordings. The input layer consists of 300 nodes, corresponding to a 10ms window. The output layer consists of 180 nodes corresponding to 2-8ms in the input window. Each output node provides a score between 0 and 100 indicating the likelihood that the corresponding input frame contains a waveform trough. The 4 convolutional layers have a kernel size of 31 frames and a stride of 1 frame so that each output node makes a prediction based on the signal in 2ms before until 2ms after the corresponding frame in the input. B) Example of averaged waveform footprint detected by Kilosort2. The purple traces were selected for training/validating the CNN. C) Example of training/validating sample creation and model prediction. i: A 10ms sample of recording specific noise is taken. ii: A waveform shape is selected from the training or*

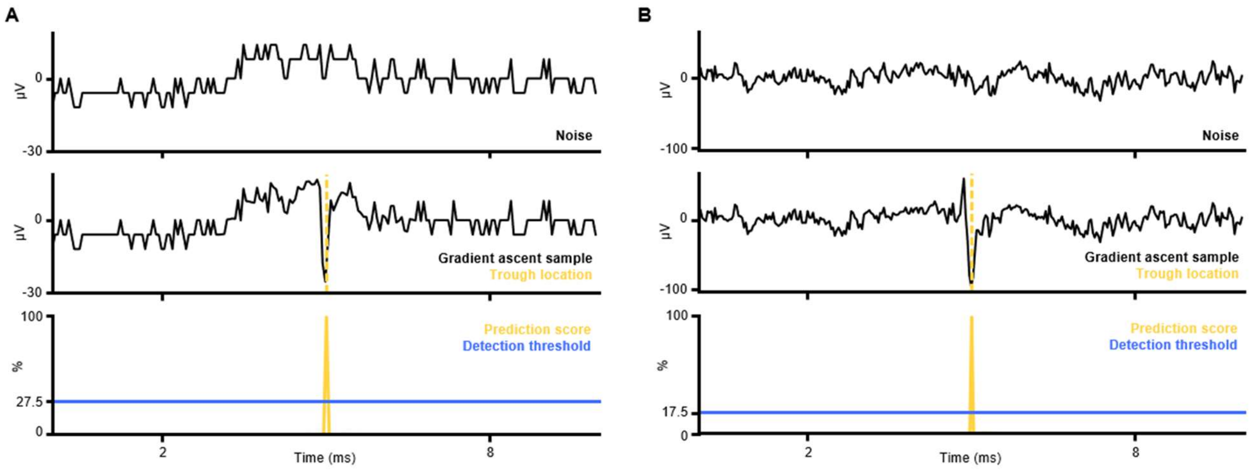


***Supplementary Figure 2: Gradient ascent forms genuine looking waveform shapes from recording noise.*** *A) (top) Intrinsic noise from an MEA electrode. (middle) Same piece of intrinsic noise after using the MEA spike detection model to perform gradient ascent on the noise to generate a waveform shape that leads to a detection with 100% certainty at the center of the noise sample. (bottom) Prediction scores for the noise sample after completing gradient ascent. B) Same as A but for Neuropixels noise using the Neuropixels spike detection model.*

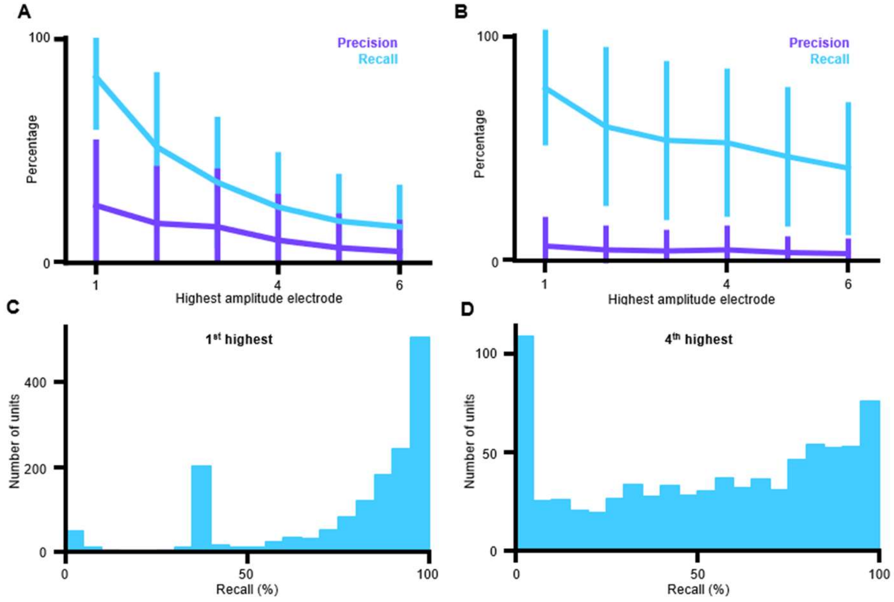


***Supplementary Figure 3: Precision and recall for Kilosort2 detections*** *A) Precision and recall of CNN spike detection model using the $i^{th}$ highest amplitude electrode for each Kilosort2 detection as ground truth. The Kilosort2 detections for 6 organoid MEA recordings are grouped together and the spike detection model is used to detect spikes on the highest amplitude electrode (1 on x-axis) down to the $6^{th}$ highest amplitude electrode (6 on x-axis). The line indicates the mean and the error bars the standard deviation over all units from all different recordings. B) Same as A but for the 6 mouse in vivo Neuropixels recordings. C) Recall distribution over all highest amplitude electrodes from the Neuropixels recordings. D) Recall distribution over all the $4^{th}$ highest amplitude electrodes from the Neuropixels recordings.*

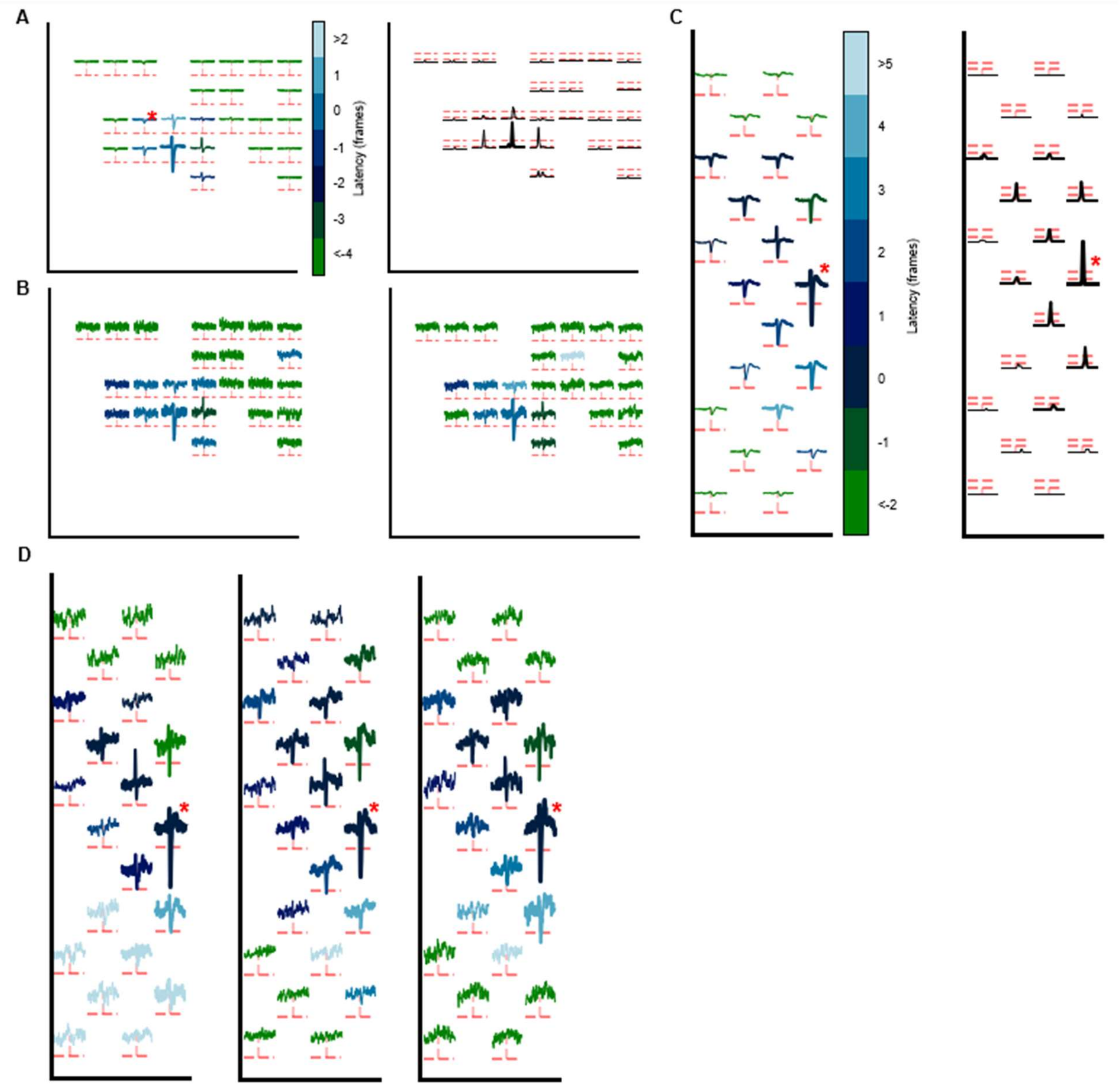


***Supplementary Figure 4: Examples of detected units with varying spike waveforms.*** *A) (left) Averaged waveform footprint of unit recorded with patch-clamp and MEA and detected by RT-Sort. (right) Corresponding averaged spike detection model footprint. Lines, colors and markers have the same meaning as Fig. 2AB. B) Examples of single spikes from the detected unit in A using RT-Sort in online mode. Color scale is the same as in A. Lines, colors and markers have the same meaning as Fig. 2D. C) (left) Waveform footprint of unit recorded with Neuropixels probe and detected by RT-Sort. (right) corresponding averaged spike detection footprint over all detected action potentials. Lines, colors and markers have the same meaning as Fig. 2AB. D) Examples of single spikes from the detected unit in C. Color scale and markers are the same as in panel C.*

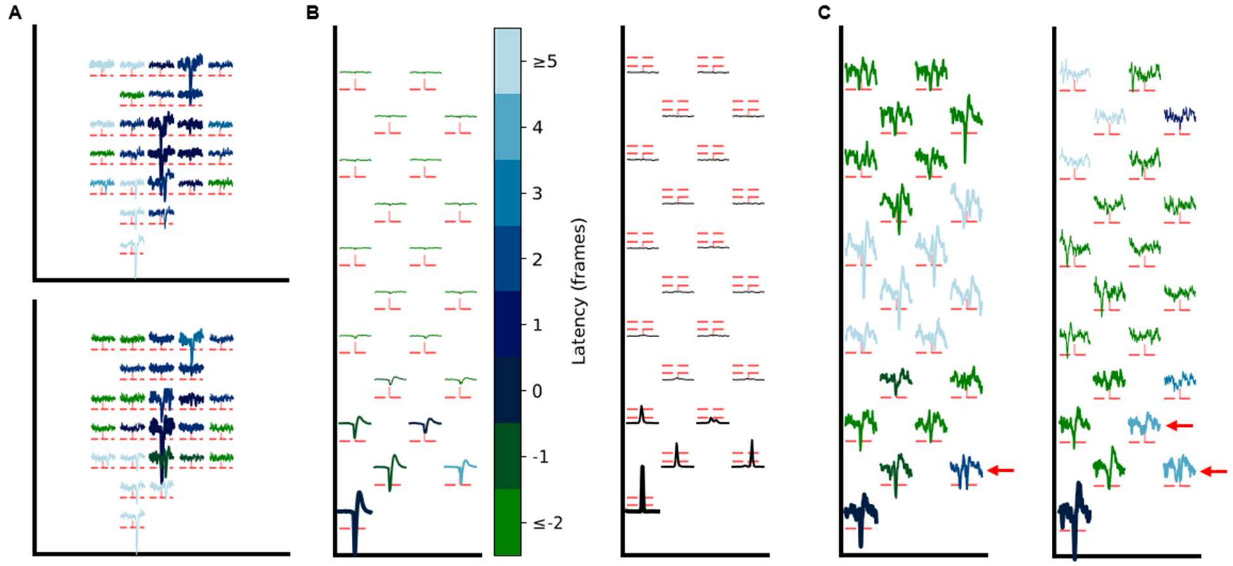


***Supplementary Figure 5: Examples of detected spikes with overlapping waveforms.*** *A) Additional examples of overlapping waveform spikes from the unit in Fig. 2A that are correctly detected by RT-Sort. Lines, colors and markers have the same meaning as in Fig. 2D. B) (left) Averaged waveform footprint of unit recorded with Neuropixels probe and detected by RT-Sort. (right) Corresponding averaged spike detection model footprint. Lines, colors and markers have the same meaning as Fig. 2AB. C) Examples of individual overlapping waveform spikes from the unit in panel B. Color scale and markers are the same as in B.*

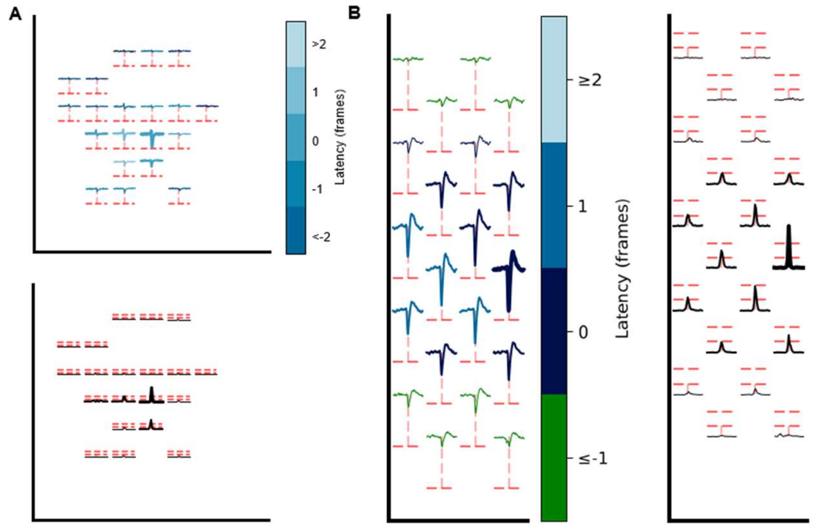


***Supplementary Figure 6: Examples of detected low amplitude units.*** *A) (top) Averaged waveform footprint of unit recorded with MEA and detected by RT-Sort. All electrodes record the unit below 5 times the signal to noise ratio, marked per electrode with the dotted red line. (bottom) Corresponding averaged CNN detection footprint. 4 electrodes detect the action potential above the loose detection threshold marked per electrode with the bottom dotted red line. Lines, colors and markers have the same meaning as Fig. 2AB. B) (left) Averaged waveform footprint of unit recorded with Neuropixels and detected by RT-Sort. All electrodes record the unit below 5 times the signal to noise ratio, marked per electrode with the dotted red line. (right) Corresponding averaged spike detection model footprint. 10 electrodes detect the action potential above the loose detection threshold marked per electrode with the bottom dotted red line. Lines, colors and markers have the same meaning as Fig. 2AB.*

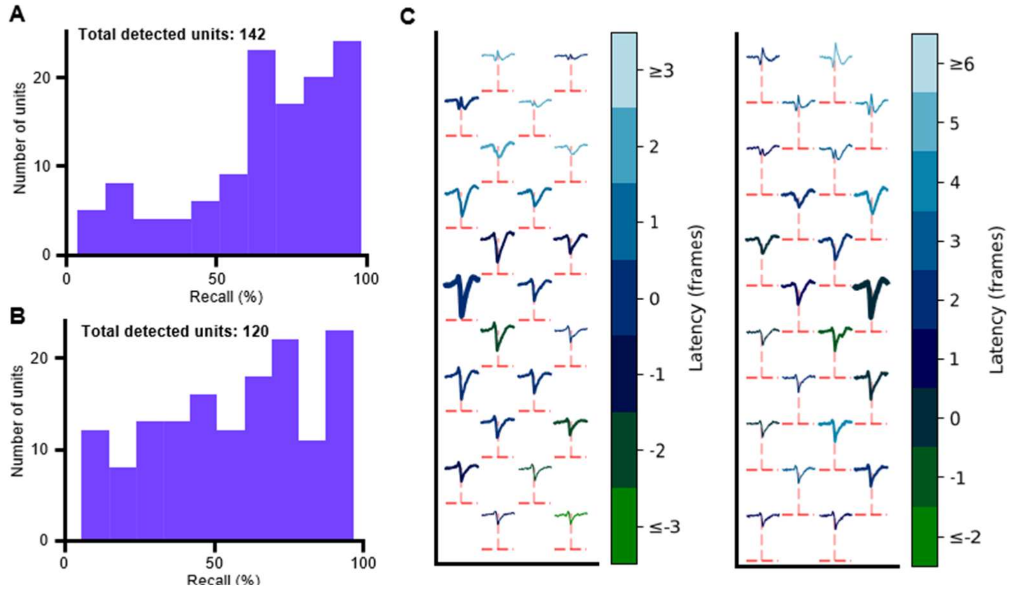


***Supplementary Figure 7: Recall on simulated ground truth recording*** *A) For each unit detected in the simulated ground truth recording using sequence metrics generated based on the ground truth spike locations, the recall over all detected spikes compared to the most similar ground truth neuron. Mean±STD = 70.9%±25.0%. B) For each unit detected in the simulated ground truth recording, the recall over all detected spikes compared to the most similar ground truth neuron. Mean±STD = 60.4%±25.9%. C) Examples of averaged waveform footprints from units from the simulated ground truth recording with unrealistic waveform shapes that did not get detected by RT-Sort (overlap score <0.01). Lines, colors and markers have the same meaning as Fig. 2D.*

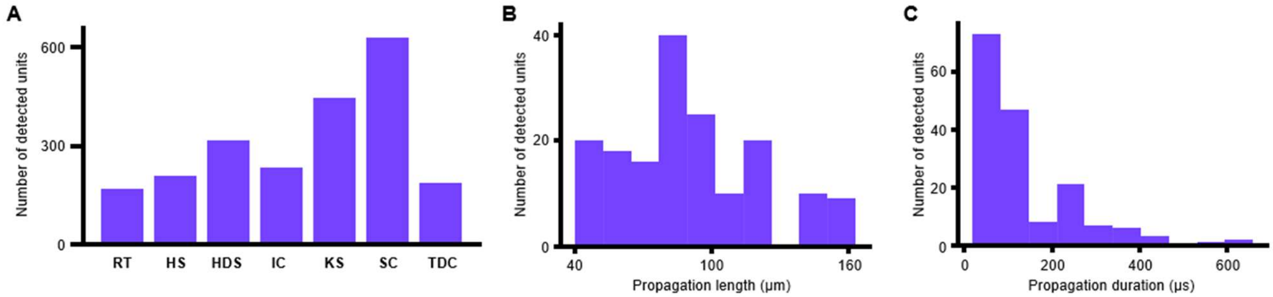


***Supplementary Figure 8: Detection statistics for a mouse in vivo Neuropixels recording.*** *A) Number of units detected by different spike sorting algorithms in the recording for Fig. 4. Abbreviations: RT = RT-Sort, HS = Herdingspikes2, HDS = HD-Sort, IC = IronClust, KS = Kilosort2, SC = SpyKing Circus, TDC = Tridesclous. B) Distribution of propagation lengths for all detected RT-Sort units in the same recording as panel A. C) Distribution of all propagation durations for all detected RT-Sort units in the same recording as panel A.*

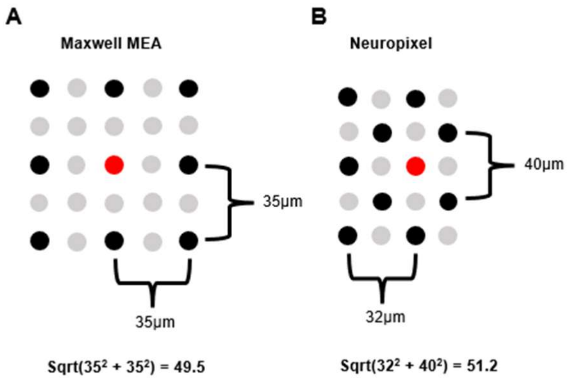


***Supplementary Figure 9: Electrode locations justify the 50µm distance threshold** A) Schematic representation of electrode locations on an MEA where every other electrode is selected in the configuration (resulting in 35µm pitch, grey electrodes are not included in the configuration). Within a 50µm radius of the electrode in red, the closest electrodes directly above and besides the red electrode are selected, as well as the closest electrodes diagonally spaced relative to the red electrode. B) Schematic representation of electrode locations on an Neuropixels probe with checkerboard configuration (grey electrodes are not included in the configuration). Within a 50µm radius of the electrode in red, the closest electrodes above and besides the red electrode are selected, as well as the closest electrodes diagonally spaced relative to the red electrode.*

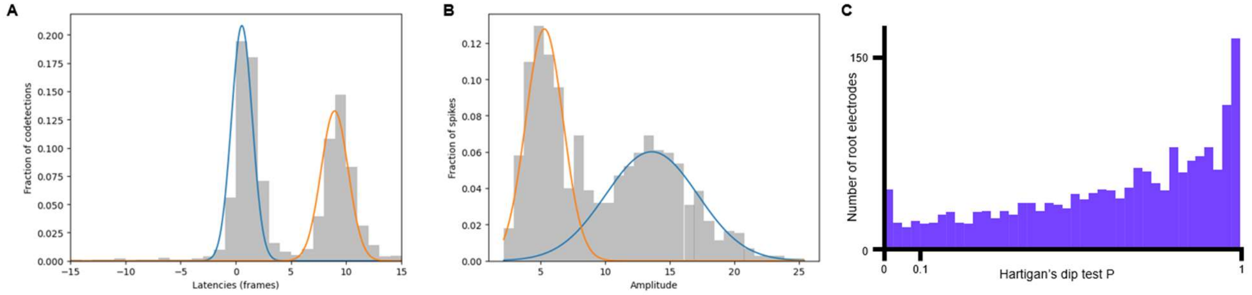


***Supplementary Figure 10: Spike cluster splitting based on amplitude and latency distributions** A) Example of an interelectrode interval distribution for all splitting codetections of a contaminated loose electrode on the Neuropixels recording of Fig. 4. The bimodal distribution reflects a propagation from two different axons. The decision boundary of the 2-component Gaussian mixture model splits the interelectrode intervals into the two separate preliminary propagations sequences. The blue and orange curves show the probability density function of the two clusters. B) Example of an amplitude distribution for all amplitudes on the root electrode of a contaminated preliminary propagation sequence on the Neuropixels recording of Fig. 4. The bimodal distribution (Hartigan's dip test, P = 0.0005) reflects a propagation from two different axons. The decision boundary of the 2-component Gaussian mixture model splits the interelectrode intervals into the two separate preliminary propagations sequences. The blue and orange curves show the probability density function of the two clusters. C) Distribution of the Hartigan's dip test P-values used for splitting the preliminary propagation sequences based on the amplitude of the root electrode in the Neuropixels recording of Fig. 4. A distribution is considered multimodal if P<0.1.*
